# BPMs regulate Arabidopsis seedling development by promoting auxin-independent degradation of the Aux/IAA protein IAA10

**DOI:** 10.1101/2024.11.26.625463

**Authors:** Zhaonan Ban, Yueh-Ju Hou, Ellyse Ku, YingLin Zhu, Yun Hu, Natalie Karadanaian, Yunde Zhao, Mark Estelle

## Abstract

After germination, seedlings undergo etiolated development (skotomorphogenesis), enabling them to grow towards the soil surface. In Arabidopsis, etiolated seedlings exhibit rapid hypocotyl elongation, apical hook formation and closed cotyledons to protect the meristem. In this study, we found that high-order mutants in the *BPM* gene family displayed defects in seedling development, characterized by a shorter hypocotyl, early apical hook opening, and opened cotyledons in the dark. BPM1, BPM2, BPM4, and BPM5 exhibit distinct expression patterns and subcellular localization in etiolated seedlings. In a hypocotyl segment assay the *bpm* mutants showed defects in auxin response indicating impaired auxin signaling in the hypocotyl. Expression of the auxin reporter *DR5:GFP* was also altered in the *bpm1,4,5* mutant in various tissues compared to the wild type. Furthermore, we showed that BPM1 and IAA10 interact in yeast two-hybrid, BiFC, and Co-IP assays. Experiments in protoplasts indicated that BPM1 promotes ubiquitylation and degradation of IAA10, and the level of IAA10 protein is greater in the *bpm1,4,5* mutant. In addition, IAA10 over-expression resulted in phenotypes similar to the *bpm* mutants. These results indicate that the BPMs target the Aux/IAA proteins for ubiquitylation and degradation. Overall, our findings shed light on the key roles of the BPMs in auxin signaling during seedling development.

## Introduction

Seedling emergence represents a pivotal phase in the plant life cycle, signifying the transition from a dormant seed to active growth. Following germination, seeds undergo two distinct developmental programs depending on whether they are growing in a dark or light environment. Etiolated development (skotomorphogenesis) occurs during seedling emergence in darkness, which is characterized in Arabidopsis by reduced root growth, elongated hypocotyls, apical hook formation, and closed cotyledons. During light-induced de-etiolation (photomorphogenesis), seedlings display reduced hypocotyl elongation, apical hook opening, along with cotyledon opening and expansion (Gendreau et al., 1997; Dong et al., 2019). Key components in light-responsive pathways and early signaling mechanisms have been identified to regulate seedling emergence. The E3 ubiquitin ligase COP1-based complex plays a central role in the light-dependent repression of photomorphogenesis in darkness; PIF (PHYTOCHROME INTERACTING FACTOR) transcription factors are also crucial contributors to this process, since *pif* high-order mutants display constitutive photomorphogenic phenotypes in darkness (Leivar et al., 2008; Shin et al., 2009; Ni et al., 2017). Beyond light signaling, phytohormones also play key roles in seedling development, such as auxin (Chapman et al., 2012; Oh et al., 2014; Du et al., 2022), gibberellic acid (Alabadi et al., 2004; Achard et al., 2007; Feng et al., 2008; Chapman et al., 2012), brassinosteroid (Li and He, 2016; Ravindran et al., 2021), ethylene (Zhong et al., 2014; Xiong et al., 2017), and jasmonate (Zheng et al., 2017). All these signals and their crosstalk regulate seedling morphogenesis (Chapman et al., 2012; Oh et al., 2014; Zhong et al., 2014; Xiong et al., 2017).

Auxin, a class of plant hormone, serves as a fundamental regulator of numerous physiological processes crucial for plant growth and development. At the cellular level, auxin regulates these processes through the control of cell division, expansion, and differentiation. The canonical auxin signaling pathway operates through the regulation of gene transcription mediated by SCF^TIR1/AFB^-Aux/IAA-ARF nuclear signaling modules. Three core components play vital roles in this pathway: the TIR1/AFB co-receptors, Aux/IAA transcriptional repressors, and ARF (AUXIN RESPONSE FACTOR) transcription factors. When auxin levels are low, the Aux/IAA proteins bind to ARFs and recruit transcriptional co-repressors TOPLESS (TPL), thereby preventing ARF activation of auxin responsive genes. When auxin levels are high, auxin promotes the binding of TIR1/AFBs to Aux/IAAs, facilitating the ubiquitylation and degradation of the Aux/IAAs. This, in turn, relieves ARF repression and activates the expression of auxin responsive genes (Lavy and Estelle, 2016). There are 6 TIR1/AFBs, 29 Aux/IAAs, and 23 ARFs in Arabidopsis, and different modules provide the basis for diverse transcriptional outputs depending on various cellular and environmental contexts (Vernoux et al., 2011; Salehin et al., 2015). A previous study reported that auxin may regulate hypocotyl elongation by mediating the IAA3-ARF6/ARF8 module (Oh et al., 2014). During etiolated development, auxin promotes cell elongation in hypocotyl development by acid growth. Auxin induces the expression of *SAUR* genes through SCF^TIR1/AFB^ signaling, and the SAUR proteins repress PP2C.D phosphatase activity, preventing the dephosphorylation of PM H^+^-ATPase. As a result, the H^+^ pump remains in the activated state, leading to apoplast acidification and PM hyperpolarization, ultimately promoting cell expansion (Du et al., 2020). However, auxin inhibits cell elongation in apical hook development, indicating a biphasic control in etiolated seedling development, which may depend on auxin levels (Du et al., 2022). Recently, the TMK1-IAA32/IAA34 signaling pathway was reported to regulate apical hook development. The receptor-like kinase TMK1 was shown to stabilize IAA32 and IAA34, thereby mediating the inhibition of concave apical hook cells (Cao et al., 2019). In addition, auxin biosynthesis and transport also affect etiolated seedlings phenotypes (Zhao et al., 2001; Vandenbussche et al., 2010).

The BPM (BTB/POZ-MATH) proteins belong to BTB/POZ protein family (BROAD COMPLEX, TRAMTRACK, and BRIC-A-BRAC/POZ and ZINC FINGER), which are substrate adaptors of Cullin3 E3 ligases. In Arabidopsis, there are six BPM proteins, named BPM1-BPM6. BPMs feature a MATH domain within their N-terminal region responsible for binding substrates, and a BTB/POZ domain in their C-terminal region for binding CUL3a and CUL3b (Genschik et al., 2013; Ban and Estelle, 2021). BPMs interact with and regulate the turnover of various transcription factors, contributing to diverse roles in plant development (Weber and Hellmann, 2009; Lechner et al., 2011; Chen et al., 2013; Chen et al., 2015; Morimoto et al., 2017; Julian et al., 2019; Mooney et al., 2019; Chico et al., 2020; Skiljaica et al., 2020). Several studies have highlighted the crucial role of BPMs in plant hormone signaling. In Arabidopsis, all six BPMs interact with HOMEOBOX 6 (HB6), a class I homeodomain–leucine zipper transcription factor that functions as a negative regulator of ABA responses. The CUL3^BPM^ E3 ligase targets HB6 for ubiquitylation and degradation, affecting ABA responses (Lechner et al., 2011). BPM3 and BPM5 interact with multiple PP2CAs in the nucleus, promoting their ubiquitylation and turnover in an ABA-dependent manner (Julian et al., 2019). Recent findings revealed that CUL3^BPM^ targets MYC2/3/4, key transcriptional regulators of jasmonate response, for degradation. The stability of BPM3 is enhanced by jasmonate, indicating a negative feedback regulatory loop to modulate MYC levels and activities (Chico et al., 2020). Despite these findings, there is currently no evidence linking BPMs to auxin signaling.

Here, we show that high-order *bpm* mutants display defects in seedling development. Hypocotyl segment assays showed that auxin response is altered in the *bpm* mutants, and changes in the activity of the *DR5:GFP* auxin reporter in various tissues in the mutants further indicate defects in auxin signaling. Moreover, we identified physical interactions between BPM1 and IAA10 through yeast two-hybrid, BiFC and Co-IP assays. Experiments also demonstrated that BPM1 increased ubiquitylation level of IAA10 and decreased the stability of IAA10 when co-expressed in Arabidopsis protoplasts. There are also more IAA10 protein accumulation in *bpm1,4,5* mutant background. Additionally, IAA10 over-expression lines produced phenotypes similar to *bpm* mutants, suggesting that BPMs may act, at least in part, by targeting IAA10 protein for ubiquitylation and degradation. Overall, this study provides evidence that the BPMs play key roles in auxin signaling through IAA proteins.

## Results

### *bpm* mutants exhibit defects in seedling development in the dark

To explore the possible roles of the BPM proteins in seedling development, we first generated deletion mutants of four *BPM* genes (*BPM1*, *BPM2*, *BPM4*, *BPM5*) using CRISPR-Cas9 technology, and named them *bpm1-2*, *bpm2-2*, *bpm4-2*, *bpm5-2* (Supplemental Fig. S1). The four single mutants did not exhibit a visible phenotype compared to wild-type (Col-0) plants, which is consistent with a previous report (Lechner et al., 2011). To overcome possible functional redundancy, we crossed the single mutants and generated several double and triple *bpm* mutant lines. Here we simplified *bpm1-2 bpm2-2* mutant to *bpm1,2* for brevity, and applied similar abbreviations to other high-order mutants. We found that 3-day-old etiolated seedlings of the *bpm1,2* and *bpm1,4,5* lines had shorter hypocotyls and partially open apical hooks and cotyledons (Fig. 1A). We then examined other high-order mutants of these *BPM* genes and found that all mutants examined showed similar defects but to varying extents (Fig. 1, B to D). When we checked the four single mutants, only *bpm1-2* and *bpm2-2* had slightly shorter etiolated hypocotyls than wild type. No other significant differences in the single mutants compared to wild-type seedlings were observed in the dark (Supplemental Fig. S2). By analyzing the phenotypes of high-order mutants, we observed that BPM1 and BPM2 may play more important roles in hypocotyl elongation, since *bpm1,2* mutant showed the shortest hypocotyls. Regarding the apical hook phenotype, BPM4 and BPM5 may be more crucial, as the apical hooks of *bpm4,5* and *bpm2,4,5* were more open than the other lines. These results indicate that the *BPM* genes exhibit both functional redundancy and some specificity. To explore this idea further, we examined the phenotypes of *bpm1,2* at time intervals after germination in the dark (Fig. 1, E to I). The growth rate of wild-type seedlings was relatively slow up to 24 hours post germination (HPG) but then increased substantially during the 24 to 36HPG time interval. We found that *bpm1,2* exhibited a growth rate similar to wild type in the early slow stage, but then was significantly slower in the rapid phase, indicating that BPM1 and BPM2 function in the rapid elongation stage (Fig. 1, F and G). Regarding the apical hook phenotype, *bpm1,2* showed normal hook formation at 36HPG, but displayed earlier opening of the apical hook compared to wild type at later time points (Fig. 1H). The cotyledons of wild-type seedlings were closed in the dark, whereas the cotyledons of *bpm1,2* were open at later stages (Fig. 1I).

**Figure 1.**
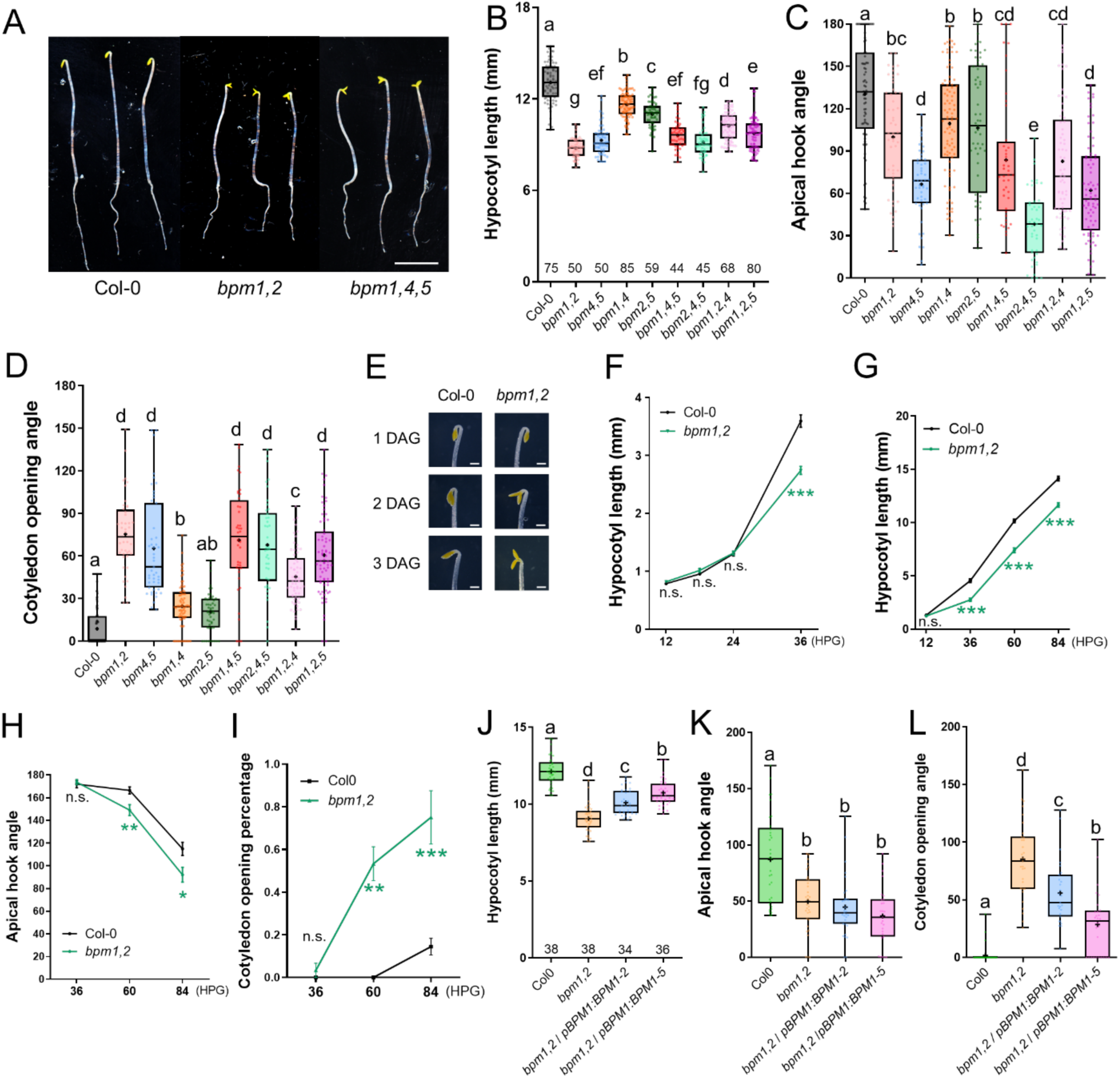
*bpm* mutants exhibit defects in seedling development in the dark. **A)** 3DAG etiolated seedlings of Col-0, *bpm1,2*, and *bpm1,4,5* mutants. Scale bar is 5 mm. **B-D)** Quantification of defects in 3DAG *bpm* and Col-0 dark-grown seedings. **(B)** hypocotyl length, **(C)** apical hook angle and **(D)** cotyledon opening angle. **E)** Closeup of apical hook and cotyledon in Col-0 and *bpm1,2* at 1-3DAG time points. Scale bar is 1 mm. **F-I)** Quantification of hypocotyl length **(F, G)**, apical hook angle **(H)**, and cotyledon opening rate **(I)** at the indicated time points during etiolated seedling development in Col-0 and *bpm1,2*. n=25 seedlings. \**P* < 0.05, \*\**P* < 0.01, \*\*\**P* < 0.001. **J-L)** Complementation of *bpm1,2* by *pBPM1:BPM1-GFP*. Quantification of hypocotyl length **(J)**, apical hook angle **(K)**, and cotyledon opening angle **(L)** in Col-0, *bpm1,2*, and two independent *bpm1,2* / *pBPM1:BPM1-GFP* lines. These data are represented as mean ± SEM. Different letters indicate statistical differences according to ordinary one-way ANOVA coupled with Holm-Sidak’s multiple comparison tests (*P* < 0.05), n.s. = not significant.

### Expression patterns of BPM1, BPM2, BPM4, and BPM5 in etiolated seedlings

Previous reports have shown that all six *BPM* genes are expressed broadly throughout the plant albeit at different levels (Lechner et al., 2011). To further explore the expression of these genes in etiolated seedlings, we generated native promoter-driven transgenic lines of *BPM1*, *BPM2*, *BPM4*, and *BPM5* fused with *GFP*. Our findings indicated that all four genes are expressed in hypocotyls, apical hooks, and cotyledons with distinct patterns (Fig. 2). *BPM1* is expressed in the elongating hypocotyl, with weaker expression in the apical hook and cotyledon. The BPM1 protein was evident in the cytoplasm and the nucleus in these tissues. It is noteworthy that *BPM1* is not expressed in the early stages of etiolated seedling growth (before 24HPG). The pattern also correlates with the elongating part of hypocotyl, with higher expression in elongating cells than in other cells (Fig. 2, A, E, I, M). The *pBPM2:BPM2-GFP* transgenic line didn’t display any fluorescence, possibly due to a low expression level. Reasoning that BPM2 may be unstable, we then treated the seedlings with the proteasome inhibitor Bortezomib. After treatment with 30 µM Bortezomib, BPM2 was clearly visible in hypocotyl and at very low levels in the apical hook and cotyledon (Fig. 2, B, F, J, N). Unlike BPM1, BPM2 is mostly localized to the nucleus. *BPM4* is highly expressed in the hypocotyl, apical hook, and cotyledon (Fig. 2, C, G, K, O), and the protein is localized to both cytoplasm and nucleus. It should be noted that there was higher accumulation of BPM4 in the cytoplasm on the concave side of apical hook. The *BPM5* gene is also highly expressed in the hypocotyl, apical hook, and cotyledon (Fig. 2, D, H, L, P). BPM5 accumulates mainly in the nucleus, with some protein in the cytoplasm of apical hook cells.

**Figure 2.**
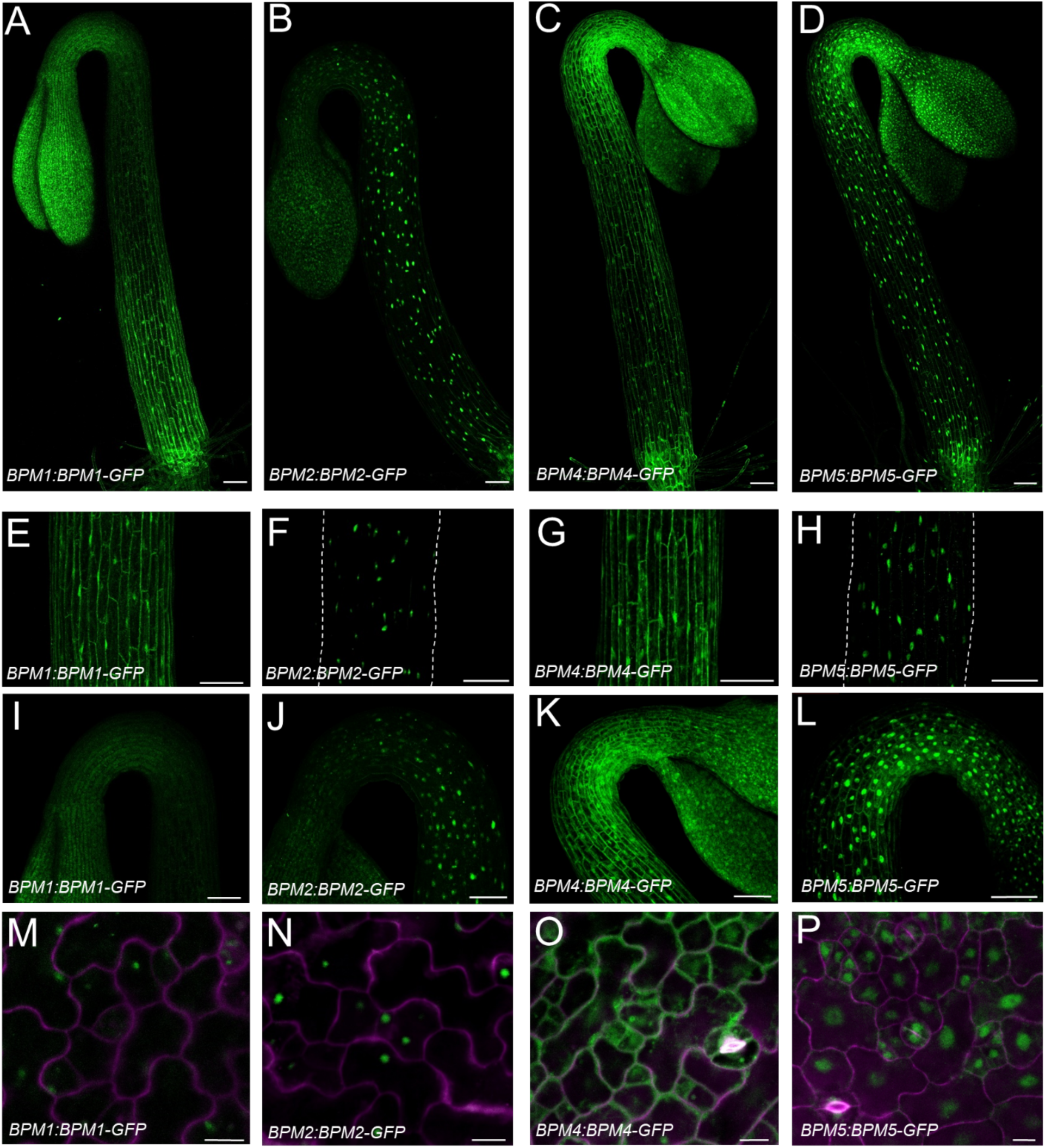
Expression pattern of *BPM1*, *BPM2*, *BPM4*, and *BPM5* genes in etiolated seedlings. GFP signal from native promoter-driven *BPM-GFP* lines showed different expression patterns. Whole etiolated seedlings **(A-D)**, hypocotyls **(E-H)**, apical hooks **(I-L)**, and cotyledon cells **(M-P)** from *pBPM1:BPM1-GFP*, *pBPM2:BPM2-GFP*, *pBPM4:BPM4-GFP*, and *pBPM5:BPM5-GFP* transgenic lines. The *pBPM2:BPM2-GFP* line was treated with 30 µM Bortezomib for 6 hours. Scale bars in **(A-L)** and **(M-P)** are 100 µm and 10 µm, respectively.

To confirm that these phenotypes were caused by loss of the *BPM* genes, we generated two independent *pBPM1:BPM1-GFP* lines in Col-0 background and crossed them with the *bpm1,2* mutant. Phenotypic analysis showed that the *pBPM1:BPM1-GFP* construct partially rescued the *bpm1,2* hypocotyl and cotyledon phenotype but not the defect in apical hook opening (Fig. 1, J to L), indicating that BPM1 plays important roles in hypocotyl and cotyledon development. We also generated two independent 35S promoter-driven overexpression lines for each *BPM* gene in Col-0 background. Phenotypic analysis showed that one *35S:BPM1* line, two *35S:BPM2* lines and one *35S:BPM5* line showed significant longer hypocotyls than wild-type seedlings in the dark, while one *35S:BPM4* line showed shorter hypocotyl than wild type, indicating that *BPM4* gene may play negative roles in hypocotyl elongation (Supplemental Fig. S3A). Regarding apical hook opening, two *35S:BPM4* lines displayed increased apical hook opening compared to Col-0 while all the other lines behaved similarly to the wild type (Supplemental Fig. S3B). None of the *35S:BPM* overexpression lines had a cotyledon phenotype. Combining all these phenotypes in the *bpm* mutants, *bpm1,2* / *pBPM1:BPM1-GFP* lines, and *35S:BPM* lines, we showed that the *BPM* genes play important roles during etiolated development in seedlings. When we examined the light-grown *bpm* seedlings, we observed that *bpm1,2* and *bpm1,4,5* mutants had slightly longer hypocotyls compared to wild-type seedlings. This phenotype became more pronounced under short-day light conditions (Supplemental Fig. S4). Since this defect was quite modest, we did not pursue further studies of light-grown seedlings.

Next, we determined whether BPM protein levels are regulated by light treatment. Western blot analysis using whole etiolated seedlings revealed that BPM4 and BPM5 levels were reduced after 6 hours of white light, while BPM1 levels did not change (Supplemental Fig. S5, A to C). As stated above, BPM2-GFP was not visible without Bortezomib treatment. In contrast, the transcript levels of *BPM1*, *BPM2*, and *BPM4* increased 1.25-fold, 1.43-fold, and 1.33-fold after light treatment compared to control, respectively, while BPM5 levels remained unchanged (Supplemental Fig. S5H). These results indicate that light exerts a post-transcriptional effect on BPM4 and BPM5 levels. We also quantified the GFP signal in the hypocotyls and apical hooks of *pBPM4:BPM4-GFP* and *pBPM5:BPM5-GFP* lines and found that the levels of BPM4 and BPM5 decreased after 6 hours of white light treatment (Supplemental Fig. S5, D and E). To explore the role of the proteasome in these changes, we treated seedlings with Bortezomib together with light and found that Bortezomib treatment increased BPM4/5 protein abundance, indicating that BPM4 and BPM5 are subject to degradation through the proteasome (Supplemental Fig. S5, F and G). These findings suggest that BPM proteins are degraded upon exposure to light, while transcript levels increase modestly.

To explore whether BPM levels are regulated by auxin, we treated *pBPM1:BPM1-GFP*, *pBPM4:BPM4-GFP*, and *pBPM5:BPM5-GFP* with 5 µM IAA for 4 hours. We found that BPM1 levels decreased by about 30% after this treatment, while BPM4 and BPM5 showed only slight decreases compared to the no-treatment control (Supplemental Fig. S6, A to C). Analysis of transcript levels of the *BPM* genes following IAA treatment showed that *BPM1*, *BPM2*, and *BPM4* levels remained unchanged, with a slight increase in *BPM5* levels. This indicates that *BPM* genes are not significantly regulated by auxin treatment (Supplemental Fig. S6D).

### *bpm* mutants have defects in auxin signaling

To assess potential defects in auxin response in the *bpm* mutants given their reduced hypocotyl length, we examined the growth response of hypocotyl segments to 5 μM IAA and NAA. Dissected hypocotyl segments from etiolated seedlings were treated with compounds, and imaged every 10 minutes for 3 hours. We found that both *bpm1,2* and *bpm1,4.5* were resistant to IAA and NAA compared to the wild type, indicating that these *BPM* genes are required for auxin-dependent growth during hypocotyl elongation (Fig. 3, A and B; Supplemental Fig. S7, A and B). We also investigated how wild type and the *bpm* mutants respond to the auxin synthesis inhibitor, L-kynurenine (Kyn). Wild-type plants exhibited reduced hypocotyl growth and a more open apical hook when treated with 10 µM Kyn. In contrast, the *bpm* mutants showed resistance to Kyn treatment in both hypocotyl and apical hook development (Fig. 3, C and D). Interestingly, in wild-type seedlings, cotyledons remained closed after Kyn treatment. In the *bpm1,2* and *bpm1,4,5* mutants, Kyn significantly inhibited the opening of cotyledons, indicating that abnormal cotyledon opening in the mutants depends on auxin (Fig. 3E). Because we observed defects in auxin response in *bpm* etiolated seedlings, we determined the transcript levels of auxin biosynthesis genes *YUCCA1, 2,* and *3* and the auxin transport genes *AUX1* and *LAX3*. We found that these genes are down-regulated in *bpm1,4,5,* with *YUCCA2/3* and *LAX3* levels at about 50% percent compared to wild type (Supplemental Fig. S8, A and B). Several *SAUR* genes, which promote cell elongation during hypocotyl elongation and regulate cell expansion during cotyledon opening, were also assessed in both hypocotyls and cotyledons. Results showed that *SAUR14*/*SAUR19*/*SAUR22*/*SAUR65* were down-regulated in hypocotyls but up-regulated in cotyledons, which suggests that expression of these genes is inhibited in hypocotyl and enhanced in cotyledon (Supplemental Fig. S8C). This is consistent with the *bpm1,4,5* phenotypes of shorter hypocotyl and opened cotyledon.

**Figure 3.**
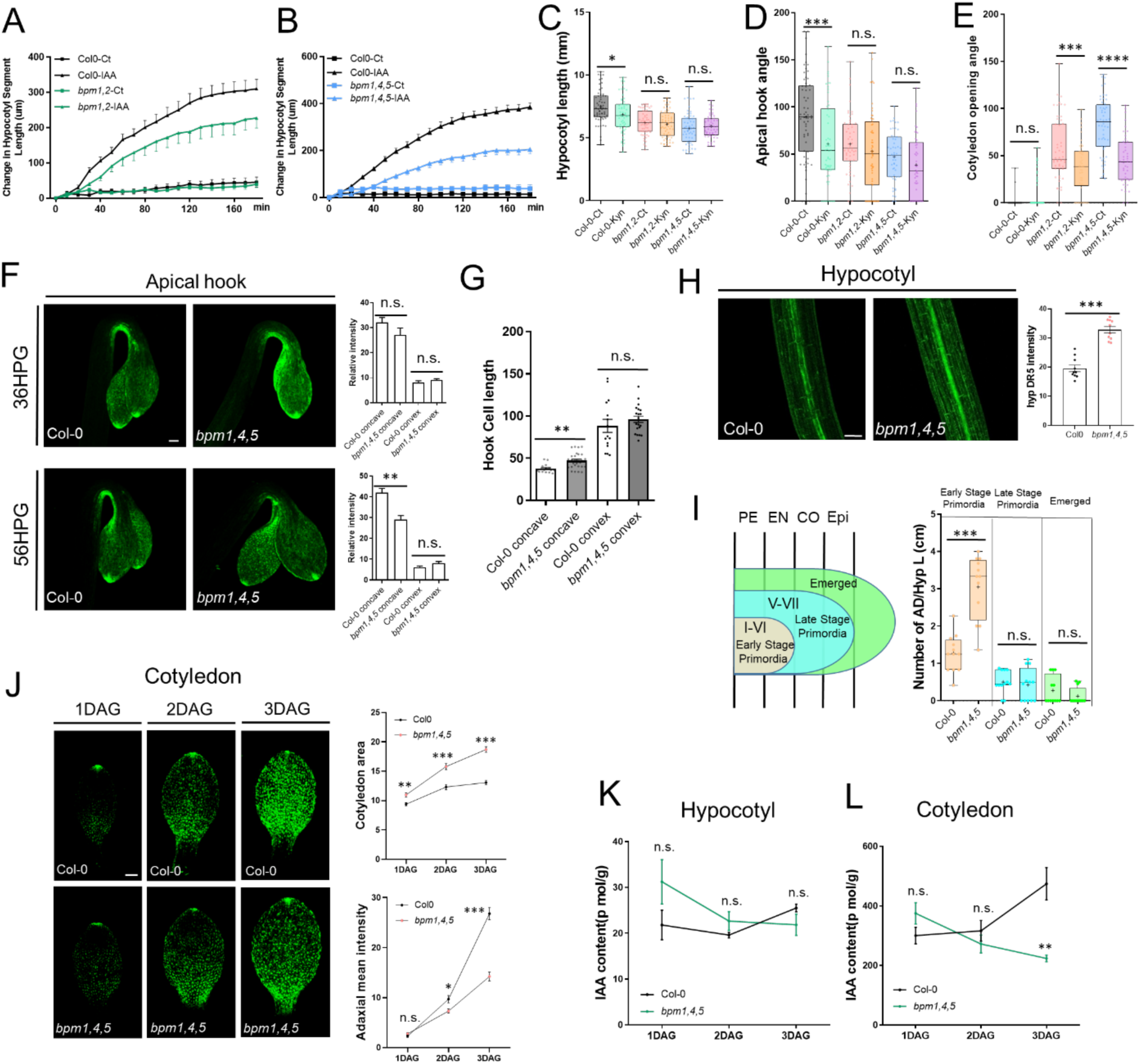
The *bpm* mutants have defects in auxin signaling. **A-B)** Hypocotyl segment elongation assay in response to 5 µM IAA for 3 hours in Col-0 and *bpm* mutants. **C-E)** Responses of Col-0, *bpm1,2*, *bpm1,4,5* etiolated seedlings to treatment with the auxin synthesis inhibitor Kyn. 10 µM Kyn was used in this assay. 2DAG hypocotyls and 3DAG apical hooks and cotyledons were used for the experiment. **F)** *DR5:GFP* signal in the apical hook region of Col-0 and *bpm1,4,5* seedlings at 36HPG and 56HPG. Quantification results are shown on the right side. Scale bar=100 µm. **G)** Cell length in apical hook regions of Col-0 and *bpm1,4,5* seedlings on concave and convex sides. **H)** *DR5:GFP* signal in 2DAG hypocotyls of Col-0 and *bpm1,4,5* seedlings. Quantification is shown on the right side. Scale bar=100 µm. **I)** Hypocotyl adventitious root primordia and emerged hypocotyl adventitious roots in Col-0 and *bpm1,4,5*. Twelve seedlings for each genotype were used. **J)** *DR5:GFP* signal on the adaxial side of cotyledons in Col-0 and *bpm1,4,5* at 1-3DAG. Scale bar=100 µm. Cotyledon area and GFP intensity were measured and shown to the right of the images. **K-L)** IAA content was measured in hypocotyls **(K)** and cotyledons **(L)** of Col-0 and *bpm1,4,5* etiolated seedlings at 1, 2, and 3DAG. These quantified data are represented as mean ± SEM.

To visualize auxin signaling in the *bpm* mutants, we generated the *bpm1,4,5 DR5:GFP* line and examined GFP signal in the hypocotyl, apical hook, and cotyledon. At 36HPG, we did not see a difference between the wild type and *bpm1,4,5* in the apical hook, while at 56HPG the GFP signal was significantly decreased on the concave side of *bpm1,4,5* plants compared to wild type, correlating with the apical hook opening phenotype of the *bpm1,4,5* mutant (Fig. 3F). We also quantified the epidermal cell length in the hook region of wild-type and *bpm1,4,5* seedlings. The cell length on the concave side of *bpm1,4,5* was longer than in wild type, while there was no difference in the convex side. These results indicate that there is reduced auxin signaling on the concave side of apical hook in *bpm1,4,5* seedlings, resulting in the failure to maintain the hook (Fig. 3G). In the hypocotyl, GFP signal accumulates in the stele region. Surprisingly, we found that the GFP signal in the stele region was stronger in *bpm1,4,5* than in wild type at 2 DAG (Fig. 3H). Subsequently, we carefully examined adventitious root development in the hypocotyl region and observed a higher number of early-stage adventitious root primordia in the hypocotyl of *bpm1,4,5* seedlings compared to the wild type. No significant differences were detected in late-stage primordia and emerged adventitious roots between *bpm1,4,5* and wild-type seedlings (Fig. 3I). Interestingly, when we examined lateral root development in light-grown seedlings, we also observed a higher number of early-stage lateral root primordia in *bpm* mutants compared to the wild type, but no differences in other stages of lateral root development (Supplemental Fig. S9). These results suggest that BPMs negatively regulate adventitious roots and lateral roots initiation. When we measured the amount of auxin in the hypocotyl of etiolated seedlings, there were no differences in auxin levels between the wild type and the *bpm1,4,5* mutant at all three time points, suggesting that the phenotype in the hypocotyl is caused by changes in auxin response (Fig. 3K).

In the cotyledon, we quantified cotyledon area and *DR5:GFP* signal on the adaxial side at 1-3DAG (Fig. 3J). At 1DAG, the area of *bpm1,4,5* was 16.9% larger than wild type. Along with cotyledon opening at 2DAG and 3DAG, the *bpm1,4,5* cotyledon area also rapidly increased, indicating rapid cell expansion. In contrast, the cotyledon area of wild type increased very slowly. There were 28.3% and 43.1% difference between *bpm1,4,5* and wild type in cotyledon area at 2DAG and 3DAG, respectively. The *DR5:GFP* signal was at a similar level at 1DAG between *bpm1,4,5* and wild type; interestingly, GFP levels increased significantly at 2DAG and 3DAG in wild type but increased very slowly in *bpm1,4,5*. We also found that IAA levels increased in the cotyledons of wild-type seedlings, but decreased in *bpm1,4,5* mutant, resulting in a significant difference in IAA levels at 3DAG between the wild type and *bpm1,4,5* line (Fig. 3L). Combined with the Kyn treatment analysis, these results suggest that cotyledon development relies on an appropriate amount of auxin, as too low or too high amounts suppress cotyledon opening and expansion.

### BPM proteins interact with IAA6/10/11 proteins

As BPM proteins function as substrate adaptors in CUL3-E3 ligases, we hypothesized that they may regulate the stability of Aux/IAA proteins, thereby modulating auxin signaling. To test this hypothesis, we used the yeast two-hybrid system to probe for interactions between BPM1 and Aux/IAAs proteins. Among the 23 Aux/IAA proteins we tested, we found that BPM1 interacts with IAA6, IAA10, and IAA11 with or without auxin supplementation, but not with the other Aux/IAA proteins (Fig. 4A). Next, we tested whether other BPMs also interact with IAA6/10/11 in the yeast two-hybrid system. We found that BPM3 and BPM4 also strongly interact with IAA6/10/11, and BPM2 has a weak interaction with IAA10, regardless of the presence of IAA. BPM5 and BPM6 do not interact with these IAA proteins (Fig. 4B). Because an interaction between IAA10 and BPM3 had been reported in an Arabidopsis interactome study (Arabidopsis Interactome Mapping, 2011), we focused on IAA10 as a representative. We further confirmed the interaction between BPM1 and IAA10 using bimolecular fluorescence complementation (BiFC) assay in *N. benthamiana* leaves and co-immunoprecipitation (Co-IP) assay in Arabidopsis mesophyll protoplasts (Fig. 4, C and D). Previous studies have shown that BPMs recognize a motif in substrates called the SBC motif (Zhuang et al., 2009; Morimoto et al., 2017) and we also identified an SBC motif in the IAA10 sequence (LSSSS) (Supplemental Fig. S11A). To investigate the function of this motif, we generated a mutant version of IAA10 with a substitution in the DII degron (GWP**S**L) and a version of IAA10 with substitutions in the SBC domain (LS**AAA**), and assessed their interactions with BPM1. Yeast two-hybrid analysis showed that IAA10 (WT) and IAA10 (DII) interact with BPM1 with or without IAA supplementation, whereas IAA10 (SBC) didn’t interact with BPM1 in the presence or absence of IAA. TIR1 interacted with IAA10 (WT) and IAA10 (SBC) only in the presence of IAA, while it did not interact with IAA10 (DII) under either condition as expected (Supplemental Fig. S11B).

**Figure 4.**
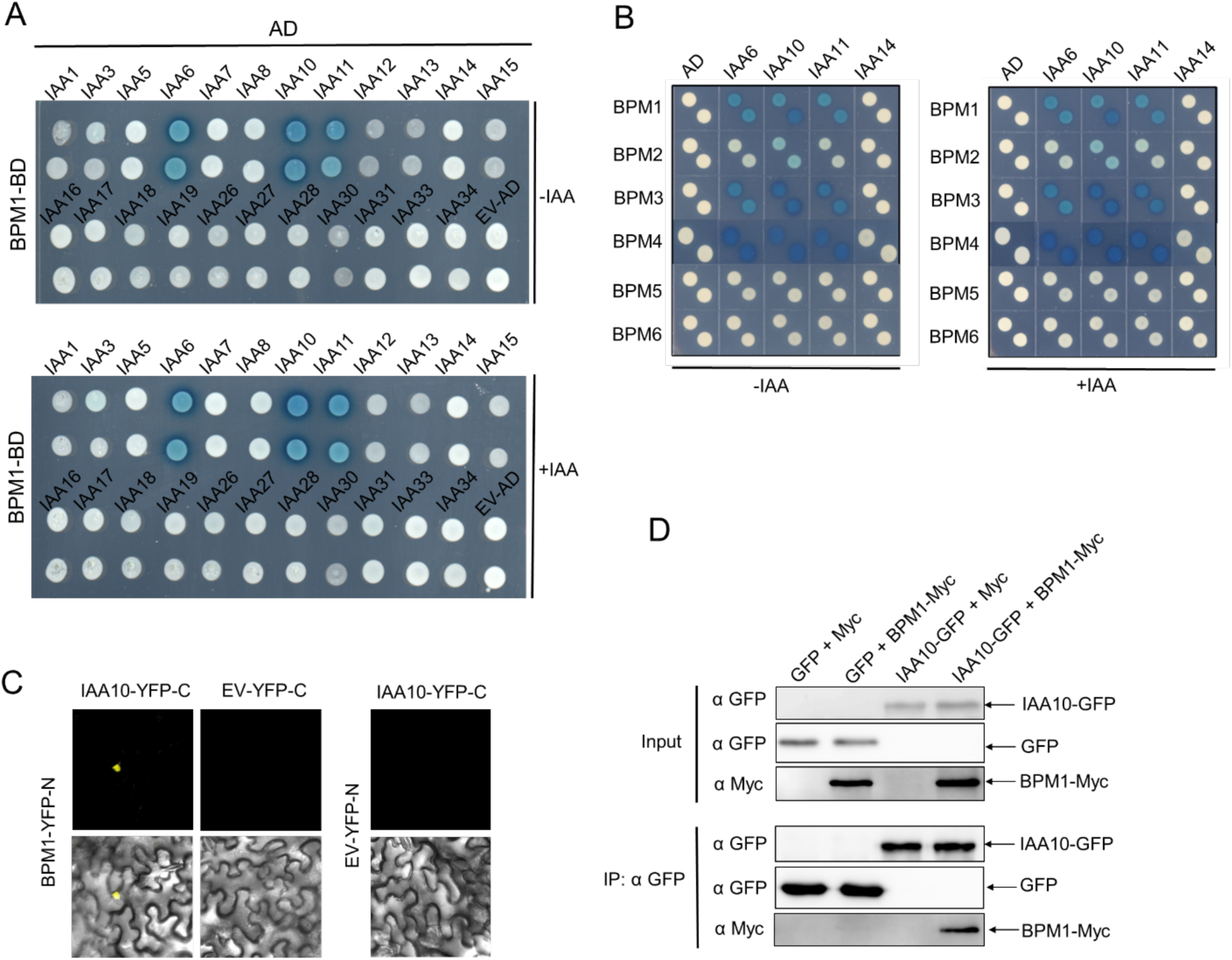
BPMs interact with Aux/IAA proteins. **A)** Yeast two-hybrid test of BPM1 with Aux/IAA proteins. **B)** Yeast two-hybrid test of six BPM proteins with IAA6/10/11/14. 50 µM IAA was used in these assays. **C)** Direct Interaction between BPM1 and IAA10 in *N. benthamiana* leaves. Confocal images of YFP fluorescence in leaves co-infiltrated with agrobacteria containing BPM1-YFP(N) and IAA10-YFP(C), BPM1-YFP(N) and EV-YFP(C), or EV-YFP(N) and IAA10-YFP(C). **D)** Co-immunoprecipitation of Myc-tagged BPM1 and GFP-tagged IAA10 using anti-Myc and anti-GFP antibodies. Total protein extracts were prepared from Arabidopsis mesophyll protoplasts transfected with plasmids expressing *GFP* and *Myc*, *GFP* and *BPM1-Myc*, *IAA10-GFP* and *Myc*, *IAA10-GFP* and *BPM1-Myc*.

### BPM1 regulates the ubiquitylation and degradation of IAA10

To further explore the impact of the BPM1 on IAA10 stability, we expressed BPM1-Myc and IAA10-Myc in Arabidopsis mesophyll protoplasts. Co-expression of BPM1-Myc together with IAA10-Myc resulted in decreased accumulation of IAA10-Myc compared to expression of IAA10-Myc alone, indicating that BPM1 regulates the level of IAA10. We also treated the protoplasts with IAA and found that the overall levels of IAA10 were reduced without BPM1 present, presumably due to TIR1/AFB-mediated degradation. However, we also found that co-expression of BPM1-Myc with IAA10-Myc resulted in a further decrease in IAA10 levels, suggesting that BPM1 regulates IAA10 levels independently of auxin (Fig. 5A). This is consistent with our observation that the interaction between BPM1 and IAA10 in the Y2H assay is not affected by auxin. We also constructed an SBC mutant version of IAA10-Myc and tested its degradation in a protoplast assay. First, we found that the IAA10 (SBC) mutant version accumulates more protein compared to the IAA10 (WT) version. When co-expressed with BPM1, the protein level of IAA10 (WT) decreased significantly, while the IAA10 (SBC) also decreased but to a lesser extent (Supplemental Fig. S11C). We then analyzed the degradation rate of IAA10 in protoplasts by treating them with cycloheximide (CHX), a protein synthesis inhibitor. Protoplasts were prepared to express *IAA10-Myc* with or without *BPM1-Myc*. The results showed that IAA10 protein was degraded over time following CHX treatment, and the degradation occurred much more rapidly when BPM1was co-expressed (Fig. 5B).

**Figure 5.**
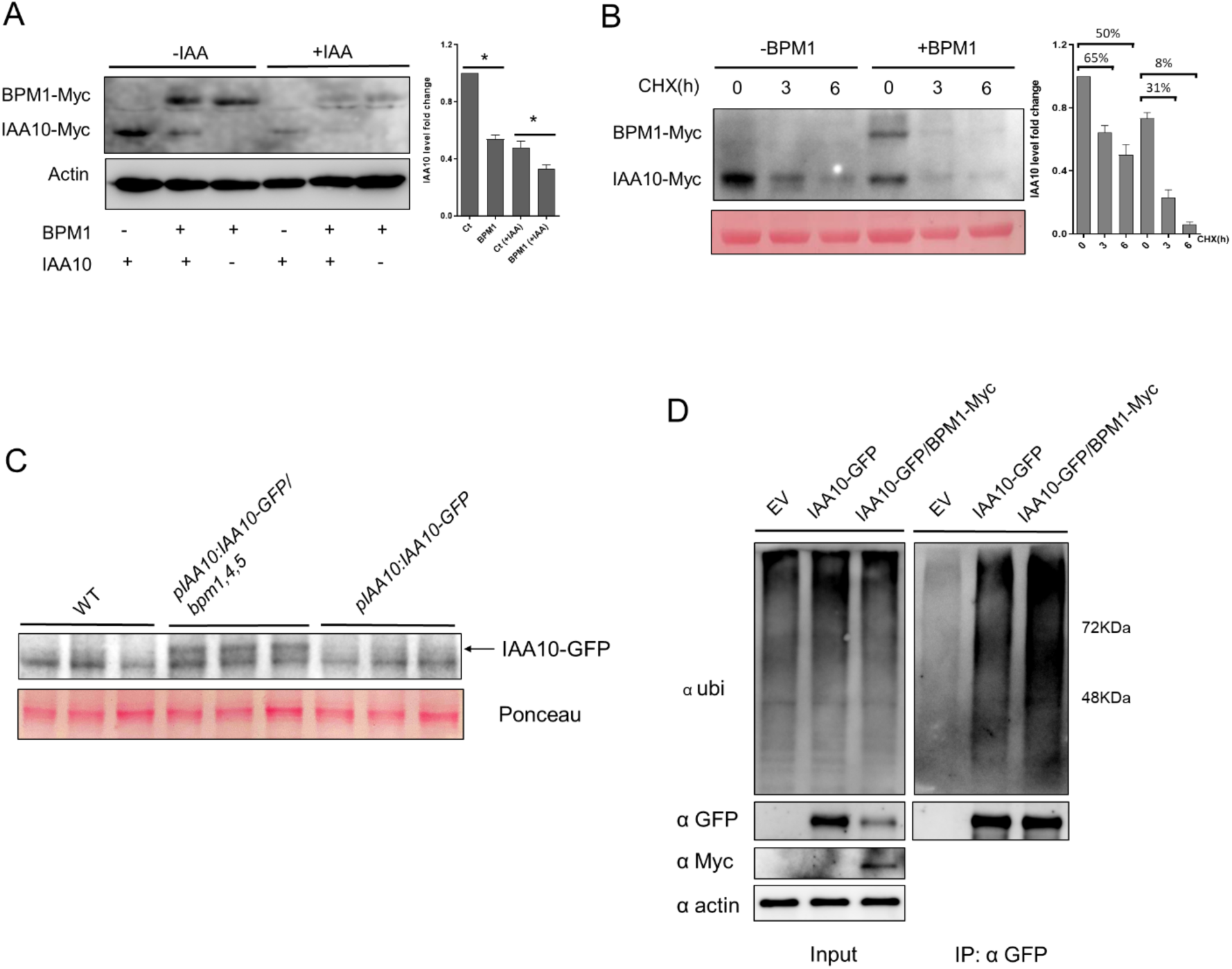
BPM1 regulates the stability of IAA10 protein. **A)** Degradation assay in Arabidopsis protoplasts. IAA10 levels were quantified with or without addition of BPM1. Protein levels of IAA10 were normalized to actin. **B)** Degradation rate of IAA10 in protoplasts. After transformation and incubation overnight, protoplasts were treated with 50 μM CHX and protein levels were analyzed at the indicated times. **C)** Western blot analysis of IAA10-GFP in *pIAA10:IAA10-GFP* and *pIAA10:IAA10-GFP/bpm1,4,5 lines*. Seedlings were treated with 5 µM IAA overnight to increase transcription. Hypocotyls of etiolated seedlings from each line were dissected for protein extraction, with three biological replicates shown in the figure. **D)** Detection of ubiquitylated IAA10 levels with or without co-expression of BPM1 in Arabidopsis protoplasts. Total protein extracts were used as input to detect overall ubiquitylated proteins, IAA10-GFP, and BPM1-Myc. Total proteins were precipitated using GFP-Trap magnetic beads, and protein loading was adjusted to obtain comparable amounts of the IAA10-GFP bands. Three biological replicates were performed in these assays.

To investigate the impact of BPMs on IAA10 protein levels *in vivo*, we initially crossed the *35S:IAA10* line with *bpm1,4,5* mutant. However, *IAA10* gene expression was silenced in the crossed line, likely due to excessive IAA10 protein accumulation. We then generated a *pIAA10:IAA10-GFP* line and crossed it with *bpm1,4,5* mutant. Given that Aux/IAA proteins are short-lived, IAA10 protein levels were very low, making detection difficult under normal conditions in both wild-type and *bpm1,4,5* backgrounds. Upon IAA treatment to enhance transcription, we observed that IAA10-GFP remained faint in the wild-type background but was detectable in the *bpm1,4,5* background, indicating increased IAA10 protein accumulation in *bpm1,4,5* (Fig. 5C). To further test whether BPM1 regulates IAA10 protein stability via ubiquitylation, we expressed *IAA10-GFP* with or without *BPM1-Myc* in protoplasts, we then used GFP-Trap beads to precipitate the IAA10-GFP protein and examined its ubiquitylated levels. Results showed that co-expression with BPM1 led to a higher level of ubiquitylation on IAA10-GFP than expression of IAA10-GFP alone, suggesting that BPM1 may target IAA10 for ubiquitylation (Fig. 5D). All together, these findings imply that BPMs function as substrate adaptors of CUL3-E3 ligase and regulate the ubiquitylation and degradation of IAA10 protein, subsequently influencing auxin signaling pathway.

### BPMs function through IAA10 protein

Since our data suggest that IAA10 is a target of the BPMs, we checked the expression pattern of *IAA10* in the *pIAA10:IAA10-GFP* line. We found that the *IAA10* gene is expressed in etiolated seedlings and the IAA10 protein accumulates in the nucleus and cytoplasm, similar to BPM1 (Fig. 6, A to C). We also generated DII, SBC, DII-SBC mutant versions of *35S:IAA10* overexpression lines and examined their phenotypes (Fig. 6, D to G). The results showed that overexpression of *IAA10 (SBC)* results in defects in hypocotyl elongation, but not apical hook. *35S:IAA10(DII)* and *35S:IAA10(DII-SBC)* lines phenotypes are similar to the *bpm1,4,5* line, with shorter hypocotyls (Fig. 6F) and apical hook defects (Fig. 6G). Some etiolated seedlings from stronger *IAA10(DII)* and *IAA10(DII-SBC)* lines also displayed partially opened cotyledons and agravitropic hypocotyls and roots. When comparing *IAA10(DII) with IAA10(DII-SBC)* lines phenotypes, they did not show very significant differences. We conducted the hypocotyl segment assay using *35S:IAA10 lines*, and found *IAA10 (DII)* and *IAA10(DII-SBC)* lines exhibited resistance to IAA treatment in hypocotyl elongation compared to wild type (Fig. 6H). These findings provide further evidence that BPMs function through IAA10.

**Figure 6.**
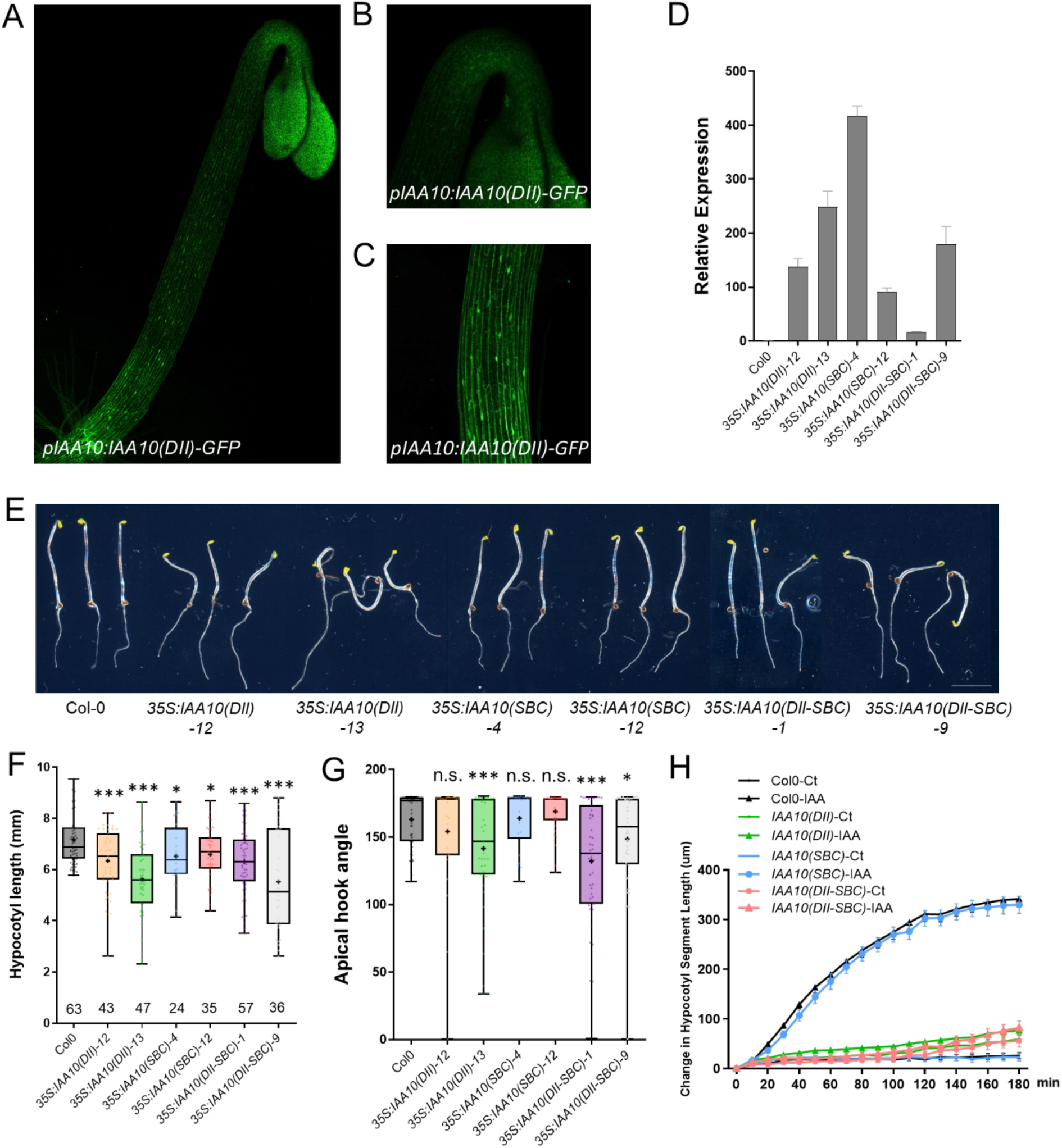
IAA10 negatively regulates hypocotyl growth and apical hook formation in the dark. **A)** GFP signal from *pIAA10:IAA10-GFP* line showed expression pattern of the *IAA10* gene. Whole etiolated seedling **(A)**, apical hook **(B)** and hypocotyl **(C)**. **D)** *IAA10* transcript levels in different versions of *35S:IAA10* lines. **E)** 2DAG etiolated seedlings of Col-0, *35S:IAA10(DII)-12*, *35S:IAA10(DII)-13*, *35S:IAA10(SBC)-4*, *35S:IAA10(SBC)-12*, *35S:IAA10(DII-SBC)-1*, and *35S:IAA10(DII-SBC)-9*. Scale bar = 5 mm. **F-G)** Quantification of hypocotyl and apical hook angles in *35S:IAA10* lines compared to Col-0. Statistical differences according to t-test analysis by comparing each genotype to Col-0. Data are represented as mean ± SEM. \**P* < 0.05, \*\**P* < 0.01, \*\*\**P* < 0.001. **H)** Hypocotyl segment elongation in response to 5 µM IAA in Col-0 and *35S:IAA10* lines.

## Discussion

Seedling emergence is a complicated process in which both light and plant hormones play vital roles. During skotomorphogenesis, auxin acts to regulate cell expansion through the TIR1/AFB-Aux/IAA pathway. Expansion growth occurs when auxin levels increase resulting in degradation of the Aux/IAA proteins and increased expression of auxin-regulated genes, including the *SAUR* genes. Although the SCF^TIR1/AFB^ E3 ligases are major regulators of Aux/IAA degradation, it is possible that other E3 ligases may also play a role. Here we show that members of the BPM family of proteins, substrate adapters for CRL3 E3 ligases, function in auxin signaling during etiolated seedling development. We also present evidence that BPMs may function by regulating stability of the Aux/IAA protein IAA10.

There are six members in the *BPM* gene family. Previous results, as well our studies, did not reveal a phenotype in the single *bpm* mutants suggesting that these genes have redundant functions. Here we use CRISPR-Cas9 technology to generate new loss-of-function alleles of *BPM1, BPM2, BPM4,* and *BPM5*. By constructing double and triple mutants, we demonstrated that these genes function in various aspects of etiolated seedling development including hypocotyl elongation, apical hook opening, and cotyledon expansion. However, individual members of the family contribute to these processes to different degrees. BPM1 and BPM2 have important roles in hypocotyl elongation, while BPM4 and BPM5 function more in apical hook maintenance. These results are also consistent with *BPM* expression patterns in different tissues. We observed *BPM1* and *BPM2* are more highly expressed in the hypocotyl region, while *BPM4* and *BPM5* are strongly expressed in the apical hook region. Interestingly, these four BPMs showed distinct subcellular localization patterns. BPM2 and BPM5 mainly accumulate in the nucleus, while BPM1 and BPM4 accumulate in both nucleus and cytoplasm. The TIR1/AFBs auxin receptors also partition differentially between the nucleus and cytoplasm. Strikingly, TIR1 is primarily expressed in nuclei while AFB1 is primarily in the cytoplasm (Prigge et al., 2020; Chen et al., 2023). Recently it was reported that cytoplasmic AFB1 functions in a non-transcriptional rapid auxin response (Prigge et al., 2020; Serre et al., 2021). In addition, AFB1 may have an inhibitory role in canonical auxin signaling (Dubey et al., 2023). It is reasonable to hypothesize that the localization of different BPM proteins may contribute to their functions. We also noticed that BPM4 may play negative roles in seedling development based on the phenotype of *35S:BPM4* lines phenotypes. It will be interesting to explore how the localization of the BPM proteins relates to their functions.

According to our results, in the dark, BPMs positively regulate cell elongation in the hypocotyl, but negatively regulate cell elongation on the concave side of the apical hook during opening. During cotyledon development, the BPMs negatively regulate cotyledon opening and expansion. Interestingly, we also found that there are more adventitious root primordia in the hypocotyls of the *bpm1,4,5* mutant, indicating BPMs negatively regulate adventitious roots formation. Similarly, BPMs negatively regulate the formation of lateral roots under light condition. Combining these observations, it appears that auxin signaling is substantially mis-regulated in the *bpm* mutants. Since we found that several BPM proteins interact with IAA10, and BPM1 regulates the stability of IAA10, it is possible that the auxin defects are related to increased levels of this Aux/IAA protein. Consistent with this, we found that overexpression of *IAA10* resulted in a phenotype similar to that of the *bpm* mutants. However, IAA10 overexpression lines did not produce a cotyledon phenotype. This defect may be due to stabilization of a different Aux/IAA protein. It is also well known that the BPMs function in a number of signaling pathways by targeting various substrates for degradation and it is possible that the defect in cotyledon development is related to a different BPM target.

Interestingly, we also found that auxin levels are altered in the cotyledons but not the hypocotyls of the *bpm* mutants. In the cotyledons of wild-type seedlings, auxin levels increase during seedling development while in the *bpm* mutants, they decrease. These results indicate that the BPMs may regulate auxin signaling, as well as auxin biosynthesis, metabolism or transport, either directly or indirectly in cotyledon. In support of this idea, we found that *YUCCA* and *LAX* genes are down-regulated in *bpm1,4,5* mutant. Because of the complexity of the auxin regulatory system, with interdependent regulation of signaling, transport, and metabolism, it is possible that the BPMs affect auxin signaling both directly and indirectly.

In the canonical auxin signaling pathway, auxin act to promote the interaction between the TIR1/AFBs and Aux/IAA proteins. In contrast, our results indicate that auxin does not contribute to the interaction between the BPMs and the Aux/IAA proteins. The TIR1/AFB proteins recognize the DII degron of Aux/IAAs and regulate their ubiquitylation and degradation. According to our results, the BPMs may recognize the SBC motif of the Aux/IAAs and regulate their ubiquitylation and degradation. BPM1 reduces IAA10 stability without IAA treatment, indicating BPMs may constitutively function in auxin pathway. We also found that BPM1 level decreased slightly with auxin treatment, while BPM4 and BPM5 were not regulated by auxin, strengthening this hypothesis. It will be very interesting and meaningful to further explore the mechanism and physiological roles of BPMs in auxin signaling pathway.

## Methods

### Plant materials and growth conditions

All Arabidopsis (*Arabidopsis thaliana*) materials used as wild-type controls and for generating transgenic lines in this study were in the Col-0 background. Seeds were sterilized with 70% ethanol for 5 minutes, washed four times with sterile water, and kept at 4°C for 5 days in darkness. Subsequently, stratified seeds were sown on half-strength Murashige and Skoog (½ MS) media plates containing 1% sucrose and 0.8% agar (Sigma-Aldrich, A1296), pH 5.7. After sowing, the seeds were exposed to white light for 12 hours to stimulate germination and then placed in continuous darkness at 22°C for the desired time for phenotypic analysis. Seedlings of Arabidopsis or *Nicotiana benthamiana* were grown in soil in growth room at 22°C under a long-day (16-hour light/8-hour dark) photoperiod. Arabidopsis plants used for protoplast preparation were grown in soil at 22°C with a short-day (8-hour light/16-hour dark) photoperiod for 4 weeks.

### Plasmid Construction and Plant Transformation

Deletion *bpm* mutants using CRISPR-Cas9 technology were generated as described previously (Gao et al., 2016). Two sgRNA sequences targeting each *BPM* gene were synthesized and amplified using *pCBC-DT1DT2* as a template. The PCR product was then cloned into *pHEE401E-mCherry* vector using HIFI DNA assembly reaction (NEB, E5520s). Since previous studies described T-DNA alleles of these genes, we designated our alleles *bpm1-2*, *bpm2-2*, *bmp4-2*, and *bpm5-2* (Julian et al., 2019; Chico et al., 2020). To construct *pBPM1:BPM1-GFP*, *pBPM2:BPM2-GFP*, *pBPM4:BPM4-GFP*, *pBPM5:BPM5-GFP* and *pIAA10:IAA10-GFP* plasmids, about 2.5 kb promoter fragments along with *BPM* and *IAA10* genomic sequences were cloned into the *pGWB4* vector. For *35S:BPM1*, *35S:BPM2*, *35S:BPM4*, and *35S:BPM5* construction, the coding sequences of *BPM* genes were cloned into the *pEarley104* vector. To construct *35S:IAA10* lines, the coding sequence of *IAA10* with DII degron mutation, SBC motif mutation and DII-SBC mutation were cloned into the *pMDC43* vector. These constructs were transformed into *Agrobacterium tumefaciens* strain GV3101, and the transformed GV3101 cells were used to generate transgenic Arabidopsis plants using the floral dip method (Clough and Bent, 1998). Transformants were selected based on their resistance to basta or hygromycin. Homozygous T3 or T4 lines were used in various experiments. The *bpm1,4,5 DR5:GFP* reporter line was generated by crossing *bpm1,4,5* with plants carrying the *DR5:GFP* transgene. Primers used for gene cloning are listed in Supplemental Table S1.

### Phenotypic characterization

Etiolated seedlings grown vertically for indicated time were used for phenotype characterization. Whole seedlings were photographed using Epson V600 flat-bed scanners. Hypocotyl lengths, hook angles and cotyledon opening angles were measured using ImageJ software. The angle of hook curvature was measured as described earlier (Zadnikova et al., 2010). Cotyledon opening angles were measured as reported previously (Li et al., 2014). Cotyledons open more than 30 degrees were counted as “opened”.

### Confocal imaging

To image BPMs-GFP, 40HPG etiolated seedlings were mounted with water and viewed with a Zeiss LSM 880 inverted microscope. Tile scan mode was used for capturing images of whole seedlings. To assess the effects of light, images from 40HPG etiolated seedlings were obtained at time 0 and after 6 hours of light with or without Bortezomib (LC Laboratories, B-1408). For IAA treatment, 40HPG etiolated seedlings were immersed in liquid ½ MS media with 5 μM IAA or control for 4 hours, then images were captured in indicated regions using the same microscope settings. To image the *DR5:GFP* lines, whole etiolated seedlings were used to capture images from apical hooks and hypocotyls at indicated time points. To assess *DR5:GFP* signal in cotyledons, cotyledons were first dissected, mounted with water, and images were captured at indicated time points.

### Hypocotyl segment elongation assay

Hypocotyl segment elongation assays were performed essentially as described earlier (Fendrych et al., 2016; Prigge et al., 2020). 40HPG etiolated seedlings were dissected using a dissecting microscope with its light source filtered with six sheets of green cello film. Roots and cotyledons were excised, and the hypocotyls were transferred to plates containing depletion medium (DM) (10 mM KCl, 1 mM MES pH 6 (KOH), 1.5% phytagel) overlain with a piece of cellophane (PaperMart.com). After 60 minutes on DM, the hypocotyl segments were transferred to auxin treatment plates (DM plus either 5 µM IAA, 5 µM NAA, or the equivalent amount of ethanol). About ten hypocotyls were used for each genotype and treatment. Using Epson V600 flat-bed scanners, the plates were scanned at 1200 dpi every 10 minutes for 3 hours. The segments were measured using a FIJI macro described earlier (Prigge et al., 2020).

### Chemical treatment

Stratified seeds were sown on ½ MS media plates supplemented with 10 µM L-kynurenine (Kyn, Sigma-Aldrich, K8625) or mock (H2O). The seeds were exposed to white light for 12 hours to stimulate germination and grown vertically for indicated time for phenotypic analysis.

### Adventitious root and lateral root quantification

Adventitious and lateral root quantification were performed using 12-day-old etiolated seedlings and 9-day-old light-grown seedlings, respectively. Seedlings were fixed as described previously (Malamy and Benfey, 1997). They were incubated in 0.24 N HCl in 20% methanol at 50°C for 15 minutes, followed by transfer to a solution of 7% NaOH and 7% hydroxylamine-HCl in 60% ethanol for another 15 minutes at room temperature. The samples were then immersed in a series of ethanol solutions (40%, 20%, and 10%) for 5 minutes each and they were stored in a solution containing 5% ethanol and 25% glycerol. The prepared samples were mounted in 50% glycerol on glass slides and examined under bright field using a 40x/1.2 NA WI objective with a DIC filter on a Zeiss LSM 880 inverted microscope.

### RNA extraction and RT-qPCR analysis

For auxin treatment, 2DAG etiolated seedlings were transferred into liquid ½ MS media with or without 5 µM IAA, then wrapped and incubated in the chamber for 4 hours. For light treatment, 2DAG etiolated seedlings were transferred into liquid ½ MS media, then wrapped or unwrapped in the growth chamber for 6 hours. Seedlings were collected and used for total RNA isolation using Qiagen RNeasy Plant Mini kit. The quality of the total RNA was assessed using a NanoDrop spectrophotometer. Two micrograms of RNA was used for RT–PCR using Thermo Maxima H Minus master mix. qRT-PCR was performed using CFX Opus 384 Real-Time PCR System (Bio-Rad) with SsoAdvanced SYBR mix (Bio-Rad). Primers used for qRT-PCR are listed in Supplemental Table S1.

### Analysis of endogenous IAA

To measure IAA levels in hypocotyls and cotyledons of etiolated seedlings, Col-0 and *bpm1,4,5* plants were sown on ½ MS media and harvested at 1DAG, 2DAG, and 3DAG. The roots were removed and the hypocotyls and cotyledons dissected and analyzed separately. The extraction, purification and the LC-MS analysis of endogenous IAA, its precursors and metabolites were carried out as describe previously (Novak et al., 2012). Briefly, approx. 5 mg of frozen material per sample was homogenized using a bead mill (27 hz, 10 minutes, 4°C; MixerMill, Retsch GmbH, Haan, Germany) and extracted in 1 mL of 50 mM sodium phosphate buffer containing 1% sodium diethyldithiocarbamate (DEDTCA) and the mixture of ^13^C6- or deuterium-labelled internal standards. After centrifugation (14 000 RPM, 15 minutes, 4°C), the supernatant was derivatized using cysteamine (0.25 M; pH 8; 1h; room temperature; Sigma-Aldrich), afterwards, the pH of sample was adjusted to 2.5 by 3 M HCl and applied on preconditioned solid-phase extraction column Oasis HLB (30 mg 1 cc, Waters Inc., Milford, MA, USA). After sample loading, the column was rinsed with 1 mL 5% methanol. Compounds of interest were then eluted with 2 mL 80% methanol. Mass spectrometry analysis and quantification were performed by an LC-MS/MS system comprising of a 1290 Infinity Binary LC System coupled to a 6495 Triple Quad LC/MS System with Jet Stream and Dual Ion Funnel technologies (Agilent Technologies, Santa Clara, CA, USA).

### Yeast two-hybrid assay

Yeast two-hybrid assays were based on the Matchmaker LexA two-hybrid system (Clontech). *BPM* genes were cloned into the *pGlida* vector. *IAA1*, *IAA3*, *IAA6*, *IAA10*, *IAA11*, *IAA13*, *IAA14*, *IAA15*, *IAA16*, *IAA17*, *IAA18*, *IAA19*, *IAA26*, *IAA27*, *IAA30*, *IAA33*, and *IAA34* genes were cloned into *pB42AD* vector by gateway LR reactions*. IAA5*, *IAA7*, *IAA8*, *IAA12*, *IAA28*, *and IAA31* genes were cloned into *pB42AD* vector by two restriction enzymes EcoRI and XhoI. Some of the Aux/IAA constructs were described previously (Calderon Villalobos et al., 2012). Constructs were co-transformed into the yeast strain EGY48, and co-transformation with empty vectors were used as negative controls. The presence of the transgenes was confirmed by growth on SD-UHW (SD/ - His/ -Trp/ -Ura) plates. Interactions were observed by detecting β-galactosidase activity on SD/Gal/Raf/X-Gal plates (with 50 µM IAA or Ethanol) after 3 days of incubation at 30°C. The primers used in this study are listed in Supplemental Table S1.

### Bimolecular fluorescence complementation assay

*BPM1-VYNE (R)* was generated by LR reaction between *pENTR-BPM1* and *VYNE(R)*, while *IAA10-VYCE(R)* was generated via LR reaction between *pENTR-IAA10* and *VYCE(R)*. A multiple cloning site was cloned into empty vectors to generate *MCS-VYNE (R)* and *MCS-VYCE (R)* as negative controls. These constructs were transformed into *Agrobacterium tumefaciens* strain GV3101 and different combinations were co-infiltrated into *N. benthamiana* leaves together with p19 strain. The transformed plants were grown in the dark for one day and then transferred to long-day conditions for 2 days. Complemented Venus fluorescence was observed using Zeiss LSM 880 inverted microscope.

### Co-IP, Degradation and ubiquitylation assays in protoplasts

For Co-IP assays, *BPM1* was cloned into *35S:Myc* plasmid and *IAA10* was cloned into *pJIT163-GFP* plasmid. Arabidopsis mesophyll protoplasts were prepared and transfected as described previously (Yoo et al., 2007). Protoplasts were transfected with *GFP/Myc*, *GFP/BPM1-Myc*, *IAA10-GFP/Myc*, or *IAA10-GFP/BPM1-Myc*, respectively, and incubated for 16 hours. Total protein from protoplast cells was extracted using protein extraction buffer (50 mM Tris-HCl (pH7.5), 150 mM NaCl, 5 mM EDTA, 1 mM PMSF, 1 mM dithiothreitol, protease inhibitor cocktail (Roche, 11836170001) and 1% Triton X-100) with 30 µM Bortezomib. After protein extraction, 20 µL of GFP-trap Agarose beads (Fisher Scientific, GTA020) was added to each reaction for 2 h at 4°C. The precipitated samples were washed three times with protein extraction buffer and then eluted by boiling the beads in SDS protein loading buffer for 5 minutes. Immunoblots were detected with an anti-GFP antibody (1:2000; Fisher Scientific, A11122), and an anti-Myc antibody (1:2000; Roche, 11814150001).

For degradation assays, *BPM1* and *IAA10* were cloned into *35S:Myc* plasmid to generate *35S:BPM1-Myc* and *35S:IAA10-Myc* constructs. An empty vector (EV) was used as a negative control. Protoplasts were transfected with *BPM1-Myc/EV*, *IAA10-Myc/EV*, or *BPM1-Myc/IAA10-Myc*, respectively, and incubated for 16 hours. Protoplasts were then treated with 5 μM indole-3-acetic acid (IAA) and incubated for an additional 4 hours. To detect the transfected proteins, total protein from protoplast cells was extracted using protein extraction buffer. Extracted proteins were separated by SDS–PAGE using 12% polyacrylamide gels, transferred to a nitrocellulose membrane and detected with anti-Myc antibodies (1:2000; Roche, 11814150001) and anti-actin antibody (1:2000; sigma-Aldrich, A0480). To examine the degradation rate, protoplasts were treated with 50 μM CHX and protein levels were analyzed at the indicated times.

In ubiquitylation assay, protoplasts were transfected with *EV/EV*, *IAA10-GFP/EV*, or *IAA10-GFP/BPM1-Myc*, respectively, and incubated for 16 hours. Protoplasts were then treated with 50 μM Bortezomib and incubated for an additional 6 hours. 500 μL total protein from protoplast cells were extracted using protein extraction buffer. Move 50 μL total protein for input, and add 25 μL equilibrated GFP-Trap magnetic beads (Bulldog Bio, GMA020) to 450 μL protein. After 2 hours rotated incubation at 4°C, the beads were gathered using a magnetic stand, washed three times with protein extraction buffer and then eluted by boiling the beads in SDS protein loading buffer for 5 minutes. Immunoblots were detected with an anti-GFP antibody (1:2000; Fisher Scientific, A11122), anti-ubiquitin antibody (Cell Signaling Tech, 3936T, 1:1000), and an anti-Myc antibody (1:2000; Roche, 11814150001).

## Author Contributions

Z.B. and Y. J-H. performed most of the experiments and analyzed the data. M.E. initiated the study. Yunde Z. designed experiments. E.K., Yinglin Z., N.K. performed experiments. Z.B. drafted the manuscript and prepared the figures, and M.E. edited the manuscript.

## Acknowledgement

Authors in the work thank Michael J. Prigge for experimental guidance. Swedish Metabolomics Centre, Umeå, Sweden (www.swedishmetabolomicscentre.se) is acknowledged for auxin analysis by LC-QqQ-MSMS. No conflict of interest is declared.

---

**Supplemental Figure S1.**
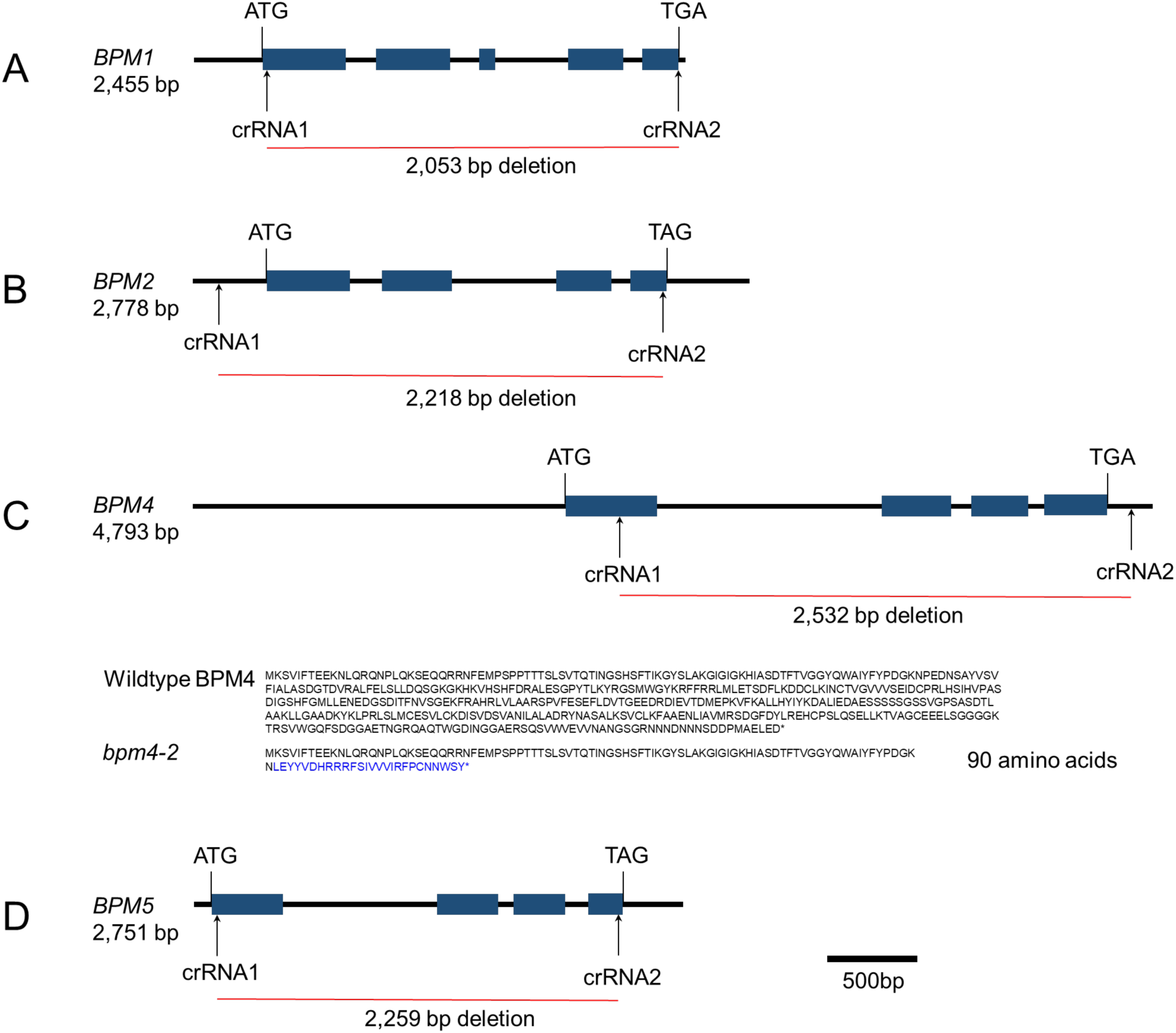
Schematic diagram of knockout positions of different *bpm* mutants. Deletion information for four *bpm* mutations created using CRISPR/Cas9 technology. **(A)** *bpm1-2*, **(B)** *bpm2-2*, **(C)** *bpm4-2*, and **(D)** *bpm5-2*.

**Supplemental Figure S2.**
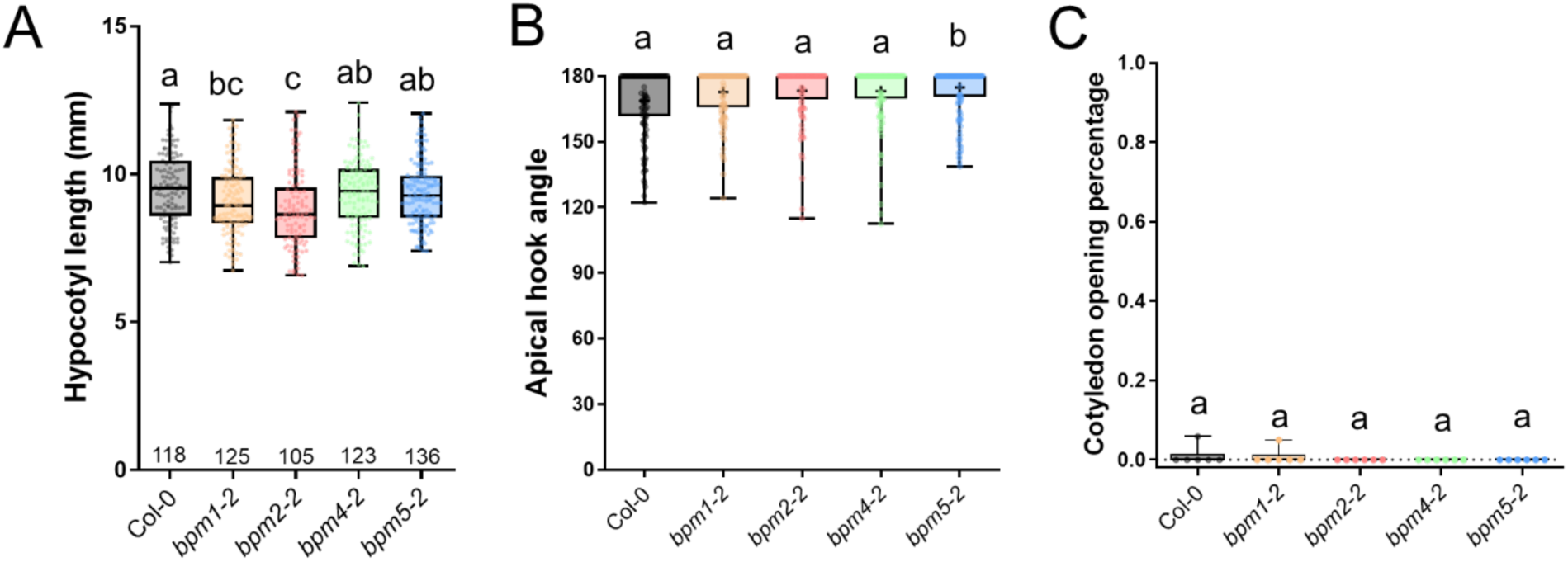
Single *bpm* mutants do not exhibit a mutant seedling phenotype. Characterization of 60HPG single *bpm* mutants compared to Col-0. **(A)** Hypocotyl length, **(B)** apical hook angle, **(C)** cotyledon opening percentage. Data are represented as mean ± SEM.

**Supplemental Figure S3.**
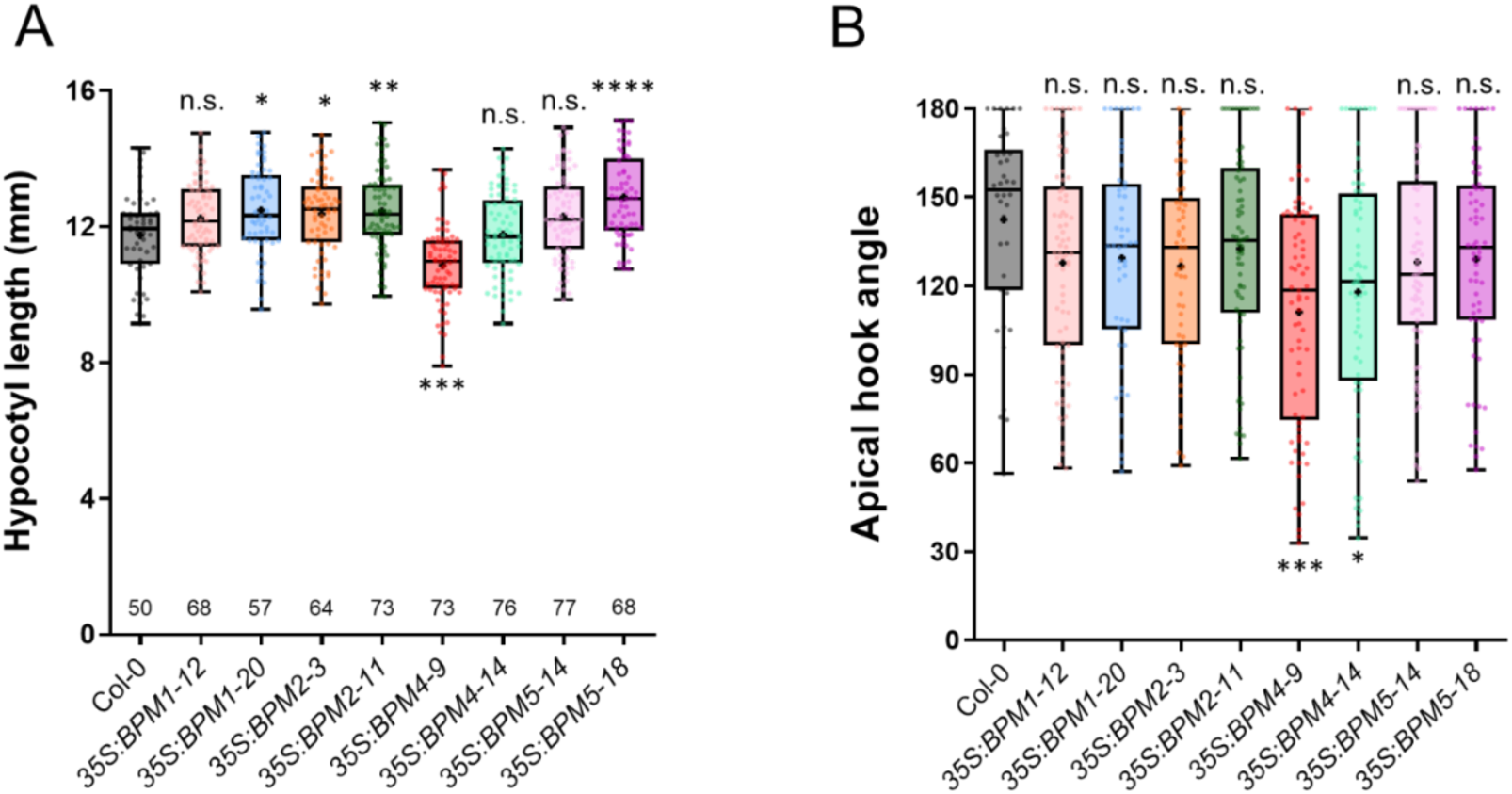
*35S:BPM* transgenic lines phenotypes. **A-B)** Analysis of 3DAG *35S:BPM* lines and Col-0. **(A)** Hypocotyl length and **(B)** apical hook angle. Statistical differences according to one-way ANOVA analysis by comparing each genotype to Col-0. Data are represented as mean ± SEM.

**Supplemental Figure S4.**
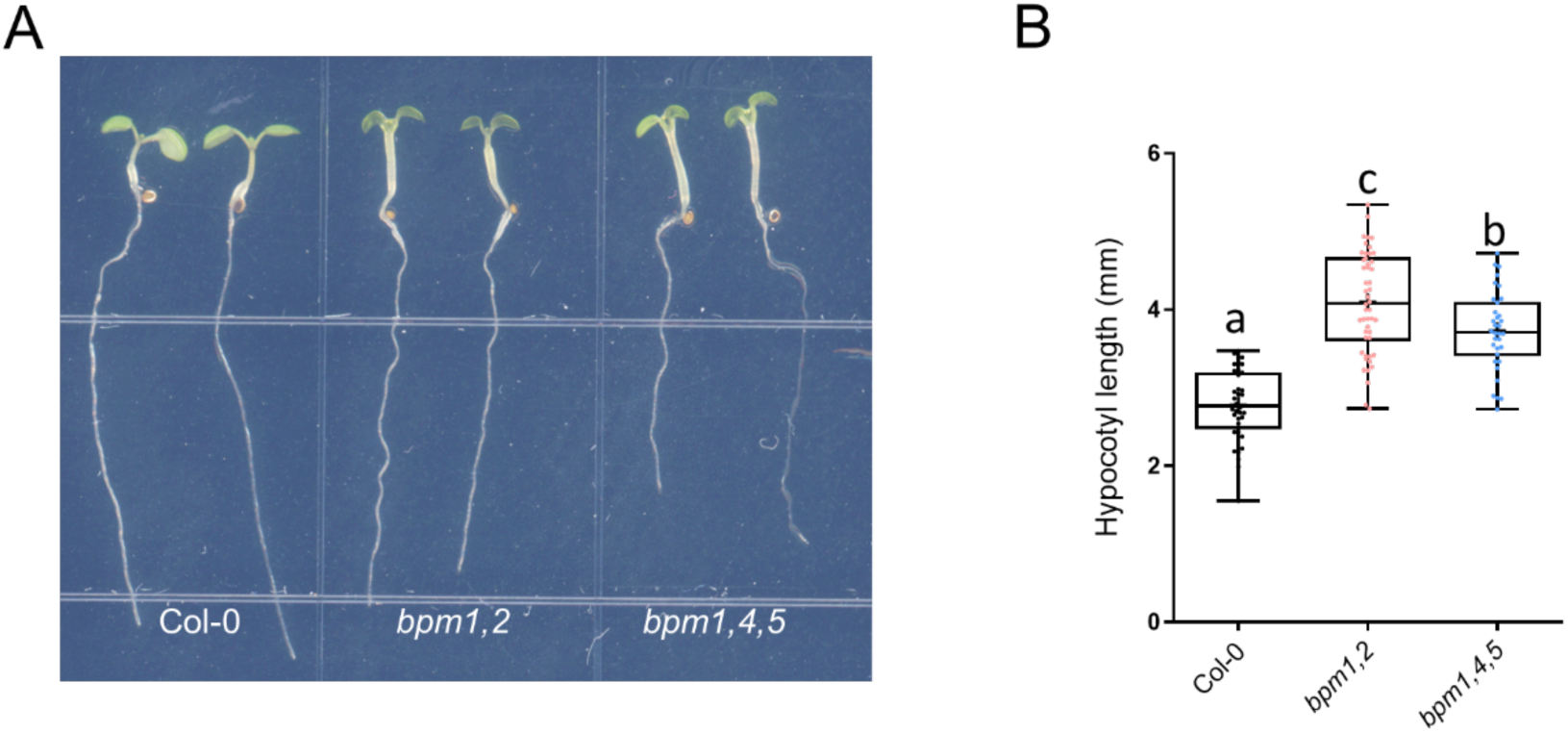
Light-grown *bpm* mutants phenotypes. **A)** 7-day old seedlings of Col-0, *bpm1,2* and *bpm1,4,5* under short-day condition. **B)** Analysis of hypocotyl length in Col-0, *bpm1,2* and *bpm1,4,5*.

**Supplemental Figure S5.**
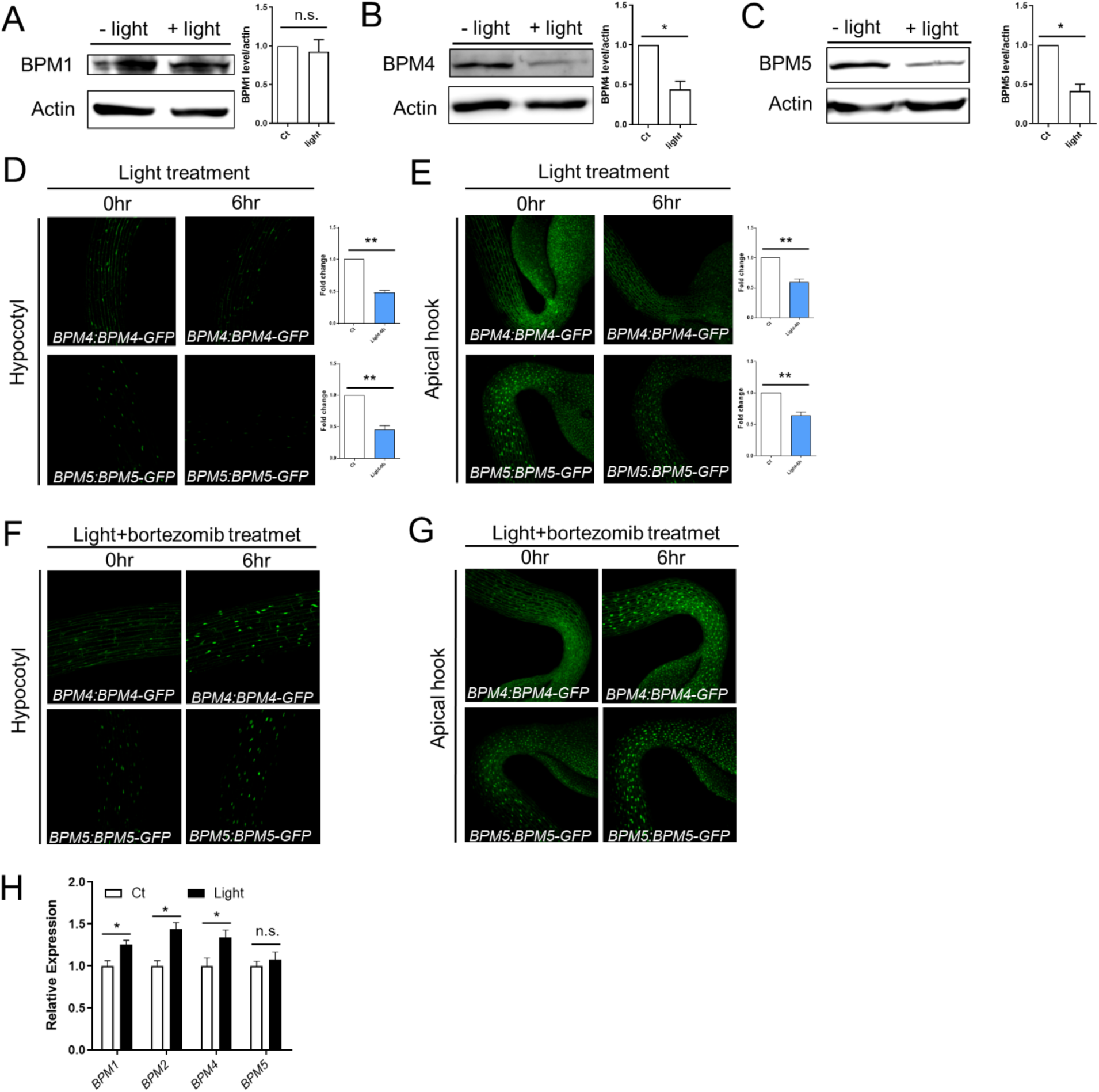
Effects of light on BPM1, BPM4, and BPM5 protein levels. **A-C)** Protein levels of BPM1-GFP **(A)**, BPM4-GFP **(B)** and BPM5-GFP **(C)** in native promoter driven transgenic plants under control (constant dark) and 6 hours white light treatment conditions. **D-E)** GFP signal in *BPM4-GFP* and *BPM5-GFP* lines after light treatment for 0 hr and 6 hours. The GFP signal in hypocotyls **(D)** and apical hooks **(E)** are shown and quantified. **F-G)** GFP signal in *BPM4-GFP* and *BPM5-GFP* lines after light plus Bortezomib treatment for 0 and 6 hours. The GFP signal in hypocotyls **(F)** and apical hooks **(G)** are shown. **H)** Transcript levels of *BPM* genes under control and light treatment conditions. Expression levels of indicated genes were normalized to *ACTIN7* expression. Light treatment expressions were presented as the fold change to the expression of control.

**Supplemental Figure S6.**
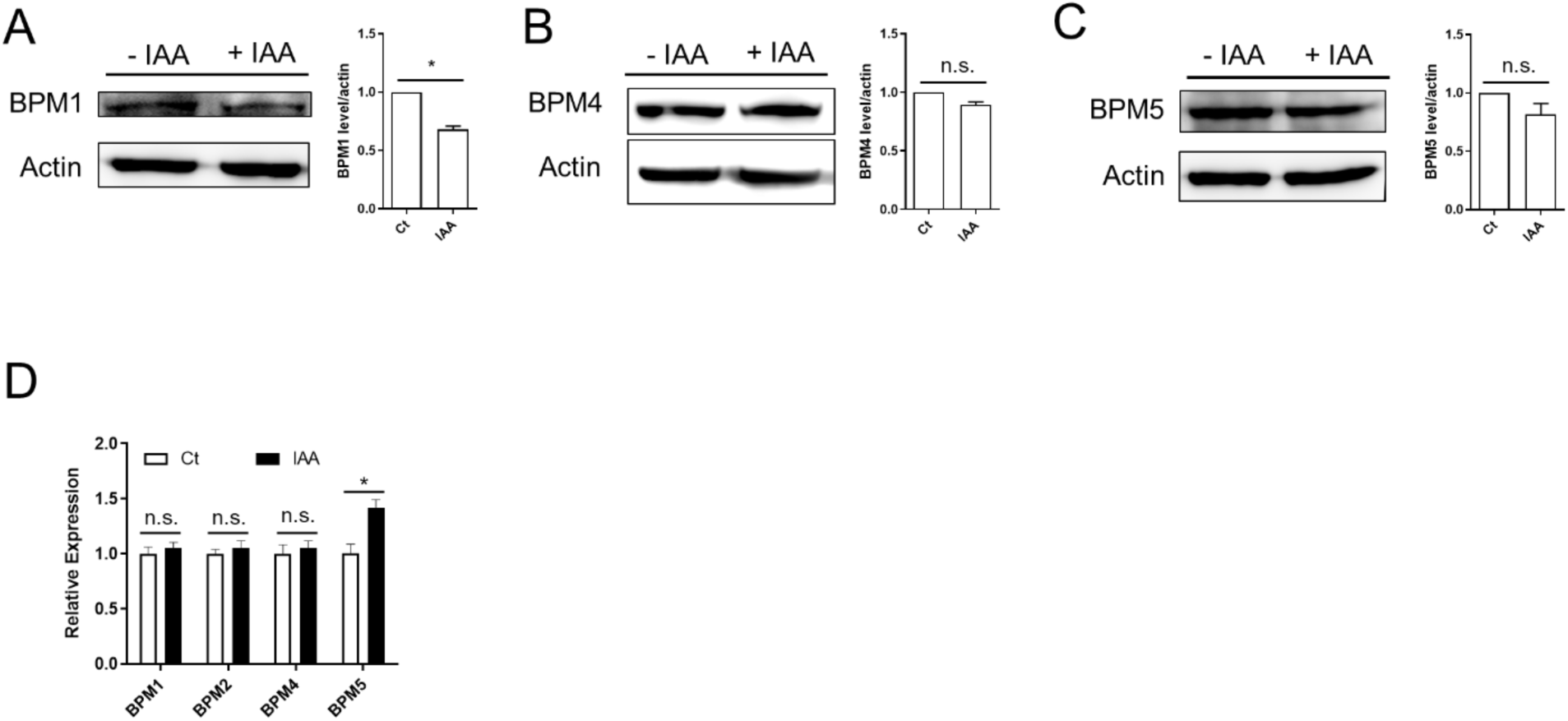
Effects of auxin on BPM1, BPM4, and BPM5 protein levels. **A-C)** Protein levels of BPM1-GFP **(A)**, BPM4-GFP **(B)** and BPM5-GFP **(C)** with or without 5 µM IAA treatment for 4 hours. **D)** *BPM* transcript levels with or without 5 µM IAA treatment conditions. Expression levels of indicated genes were normalized to *ACTIN7* expression. The effects of IAA are presented as the fold change relative to the control.

**Supplemental Figure S7.**
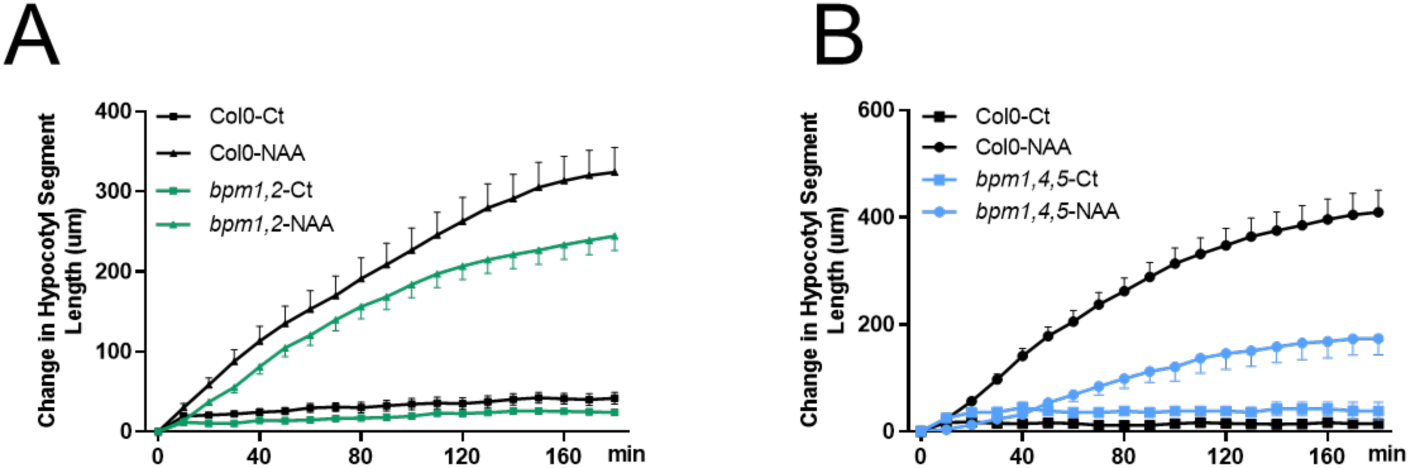
Hypocotyl segment elongation in response to NAA. Changes in hypocotyl segment length were quantified in *bpm1,2* **(A)** and *bpm1,4,5* **(B)** and compared to Col-0. 5 µM NAA was used in this assay.

**Supplemental Figure S8.**
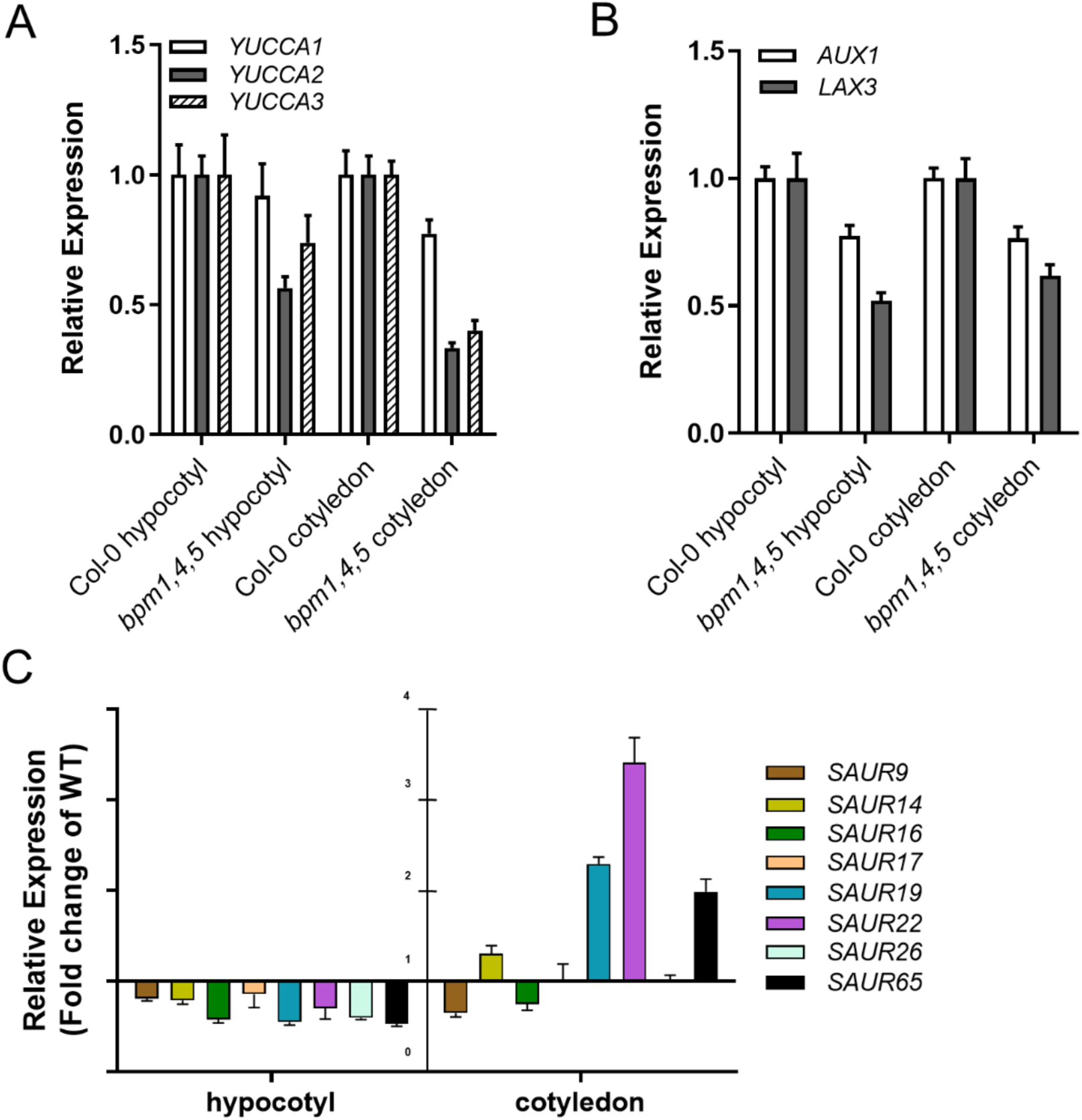
Auxin biosynthesis, transport and responsive genes transcription were affected in *bpm* mutants. **A)** Transcript levels of auxin-related genes in Col-0 and *bpm1,4,5.* **B)** Transcript levels of selected *SAUR* genes in hypocotyl and cotyledon in *bpm1,4,5.* The expression levels were presented as the fold change relative to Col-0. **C)** Expression of auxin-responsive genes in Col-0 and *bpm1,4,5* in hypocotyl segments treated with IAA.

**Supplemental Figure S9.**
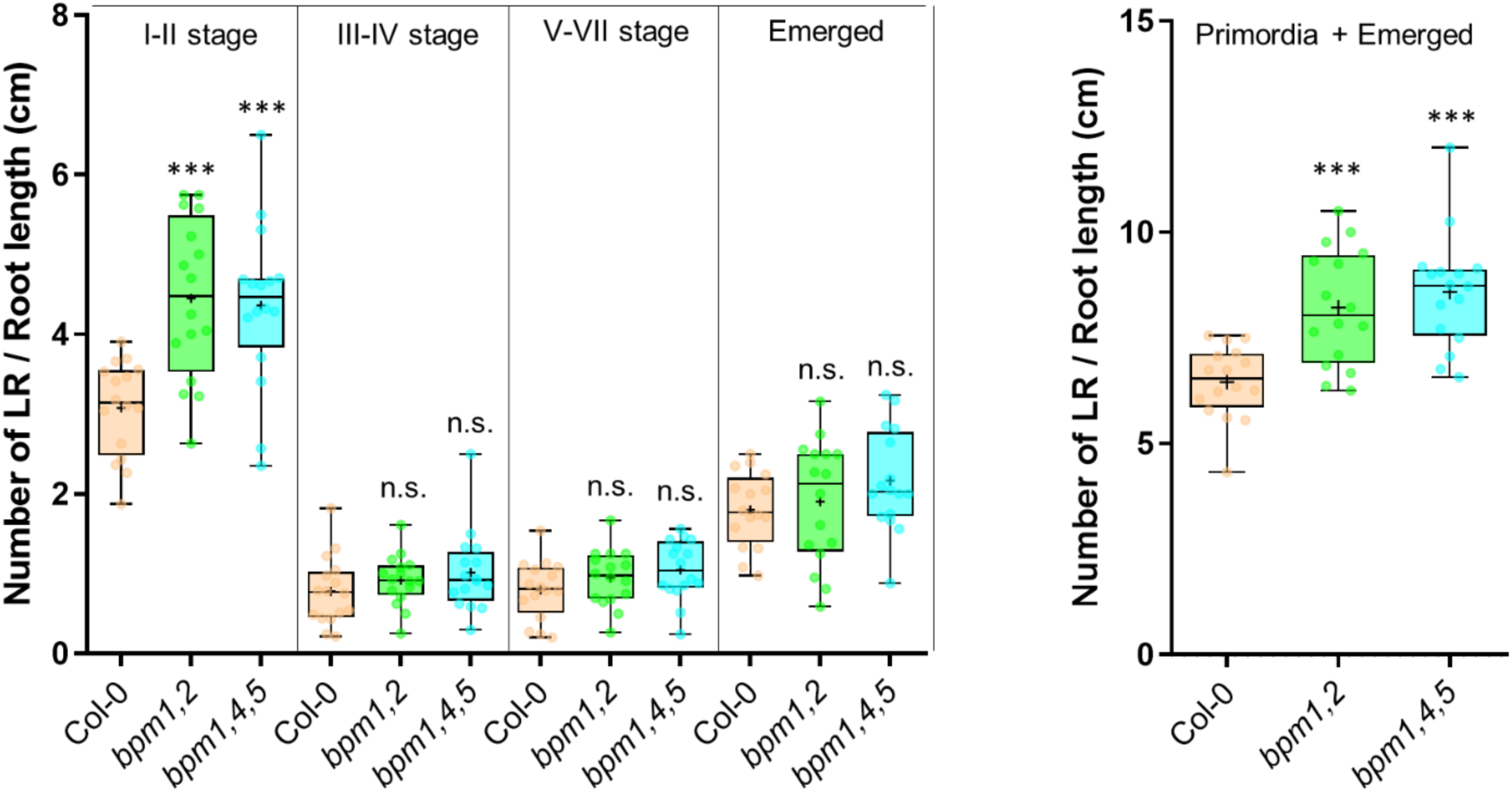
Lateral roots development in light-grown seedlings of Col-0 and *bpm* mutants. Lateral root primordia and emerged lateral roots in Col-0 and *bpm* mutants were observed in 9-day-old light-grown seedlings. Sixteen seedlings of each genotype were used for quantification.

**Supplemental Figure S10.**
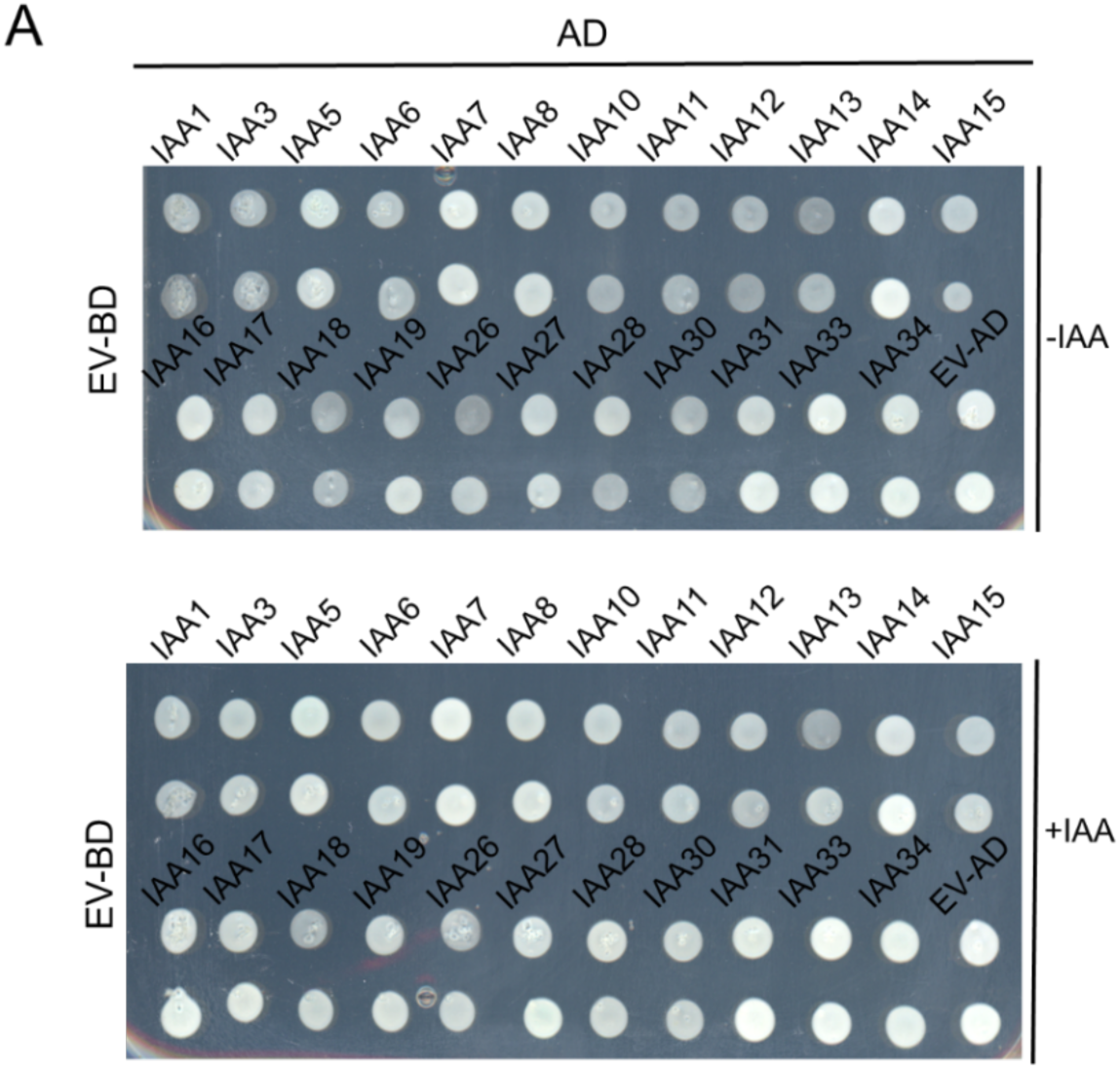
Empty vector control does not interact with Aux/IAA proteins in yeast two hybrid assay. Empty vector *pGlilda* was used in BD, and *Aux/IAA* genes were cloned in *pB42AD* vector.

**Supplemental Figure S11.**
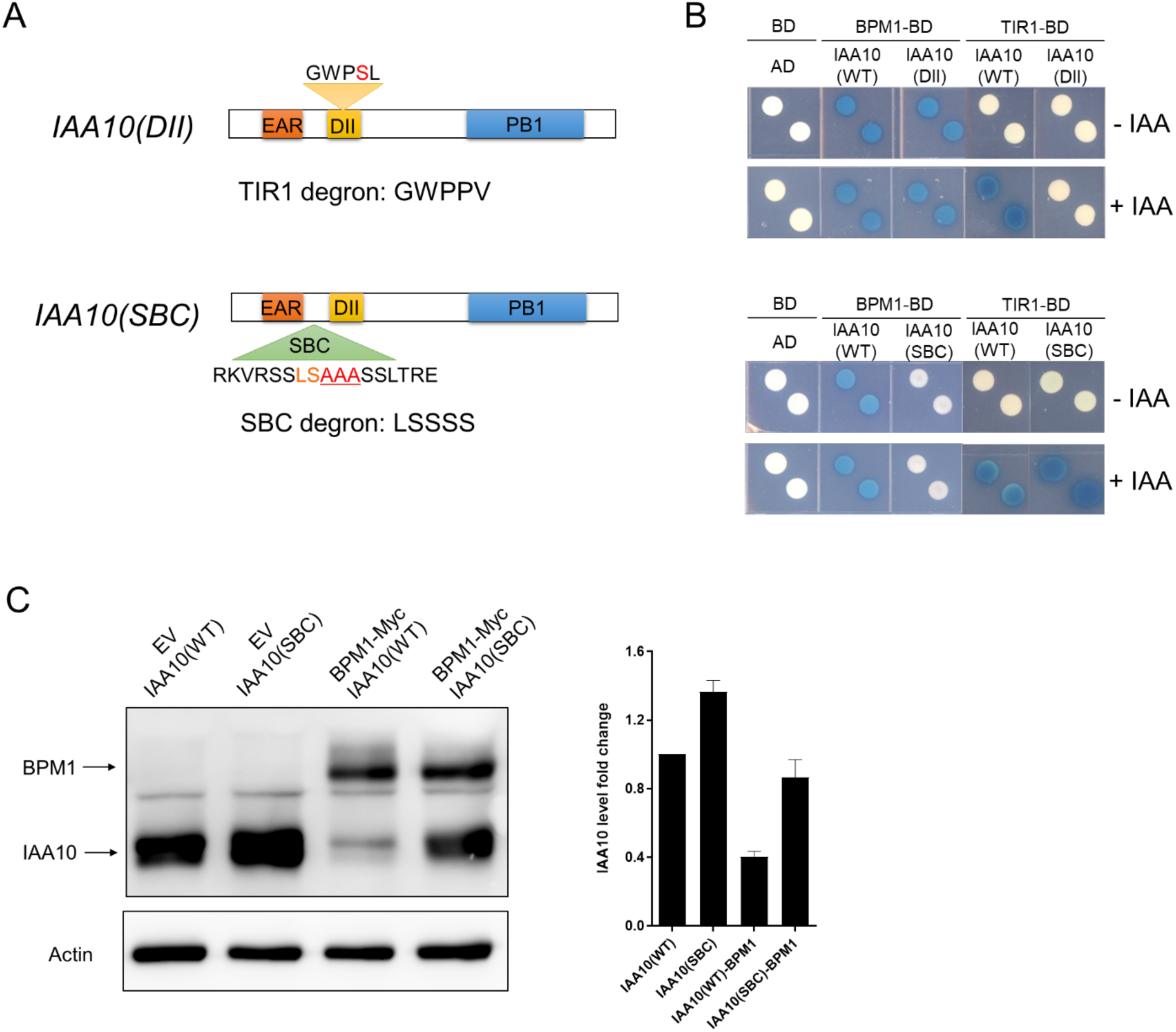
BPM1 interacts with the SBC motif in IAA10. **A)** Schematic diagram showing the positions of the DII and SBC degrons in *IAA10*. **B)** Yeast two-hybrid test of BPM1 and TIR1 interaction with different versions of IAA10. IAA10 (WT) refers to wild-type version, IAA10 (DII) stands for DII mutant version, and IAA10 (SBC) stands for SBC mutant version. **C)** Degradation assay with two versions of IAA10 (wild-type and SBC mutant versions) and BPM1 in Arabidopsis protoplasts.

**Supplemental Table S1.**
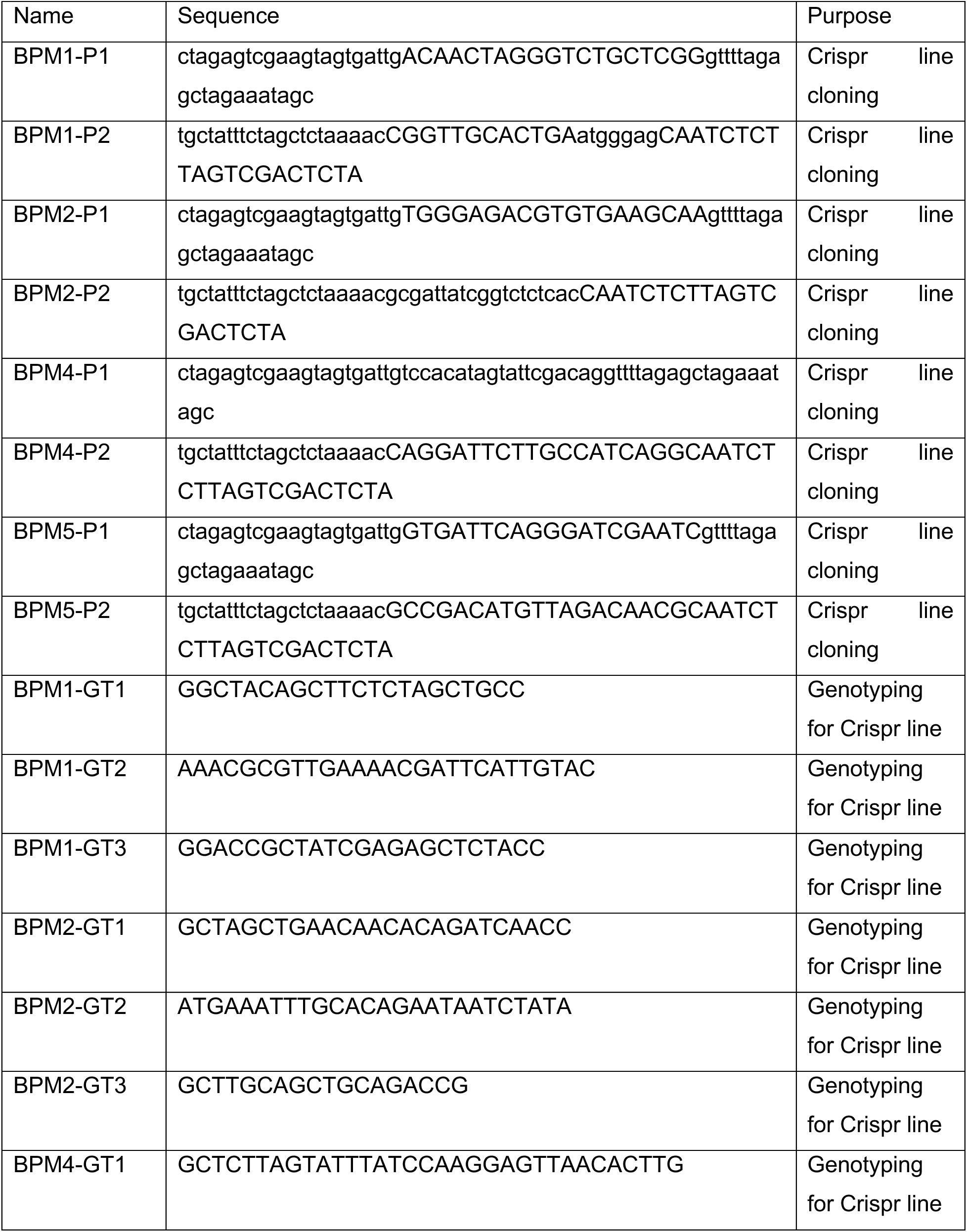
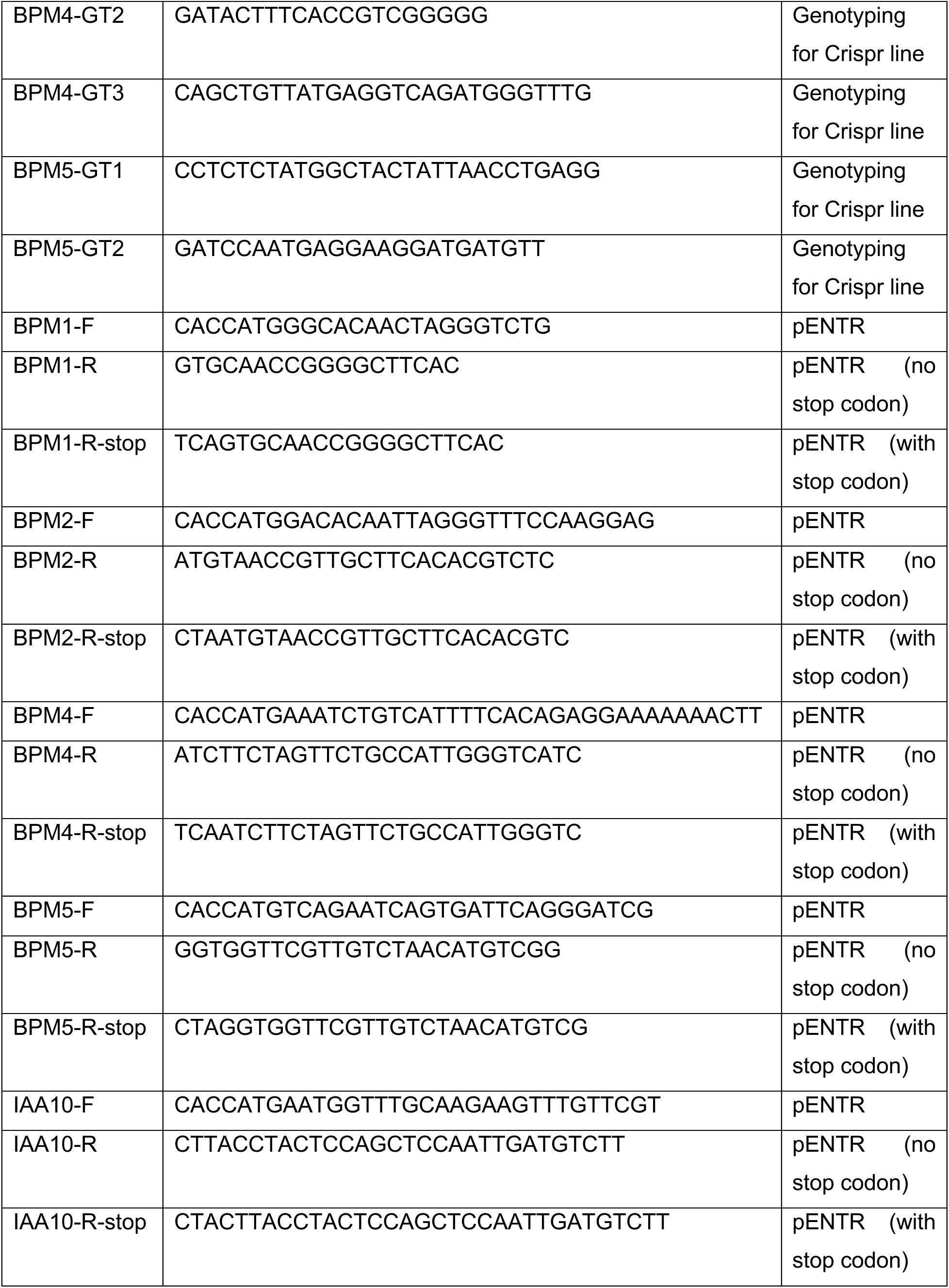
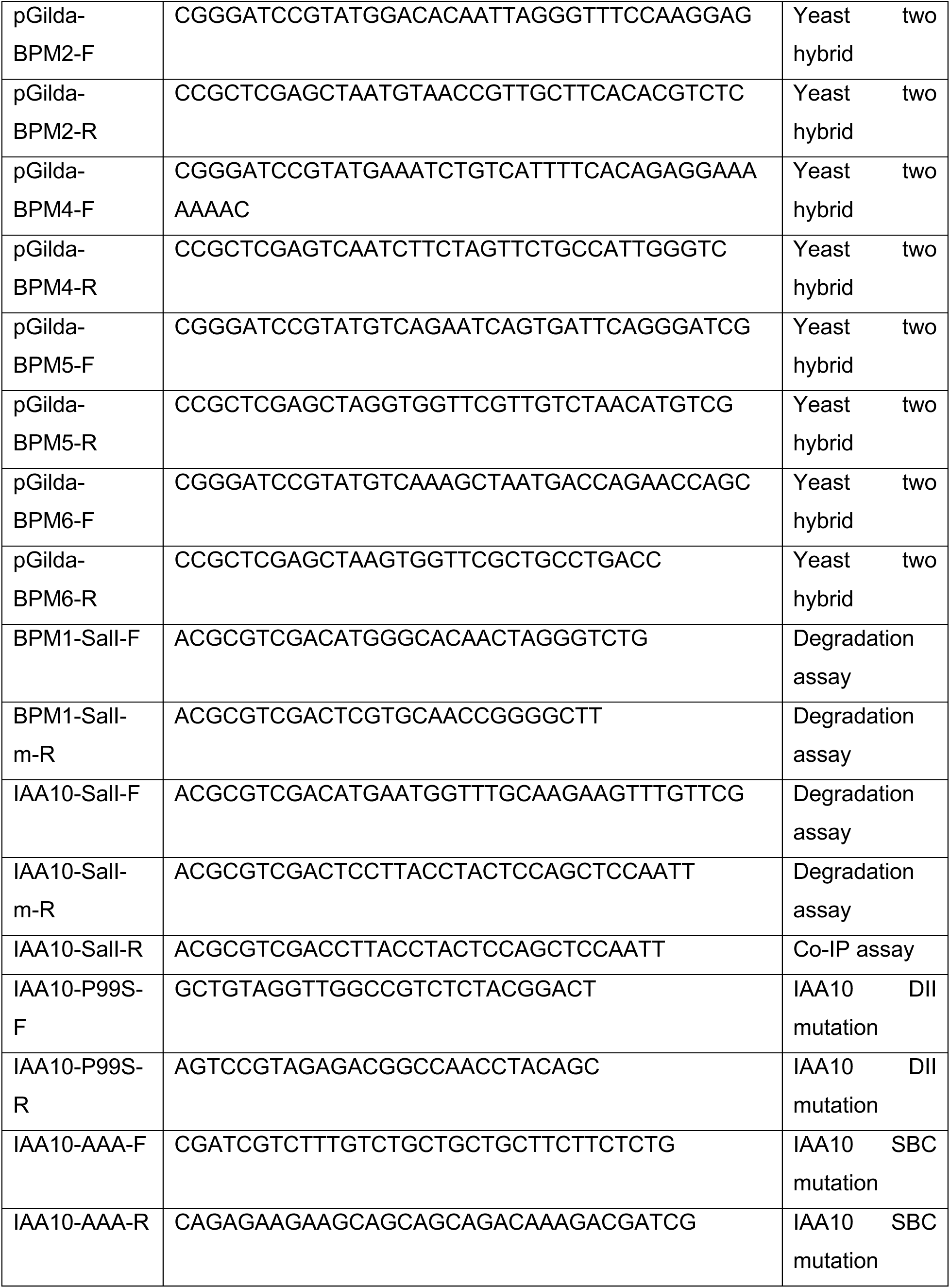
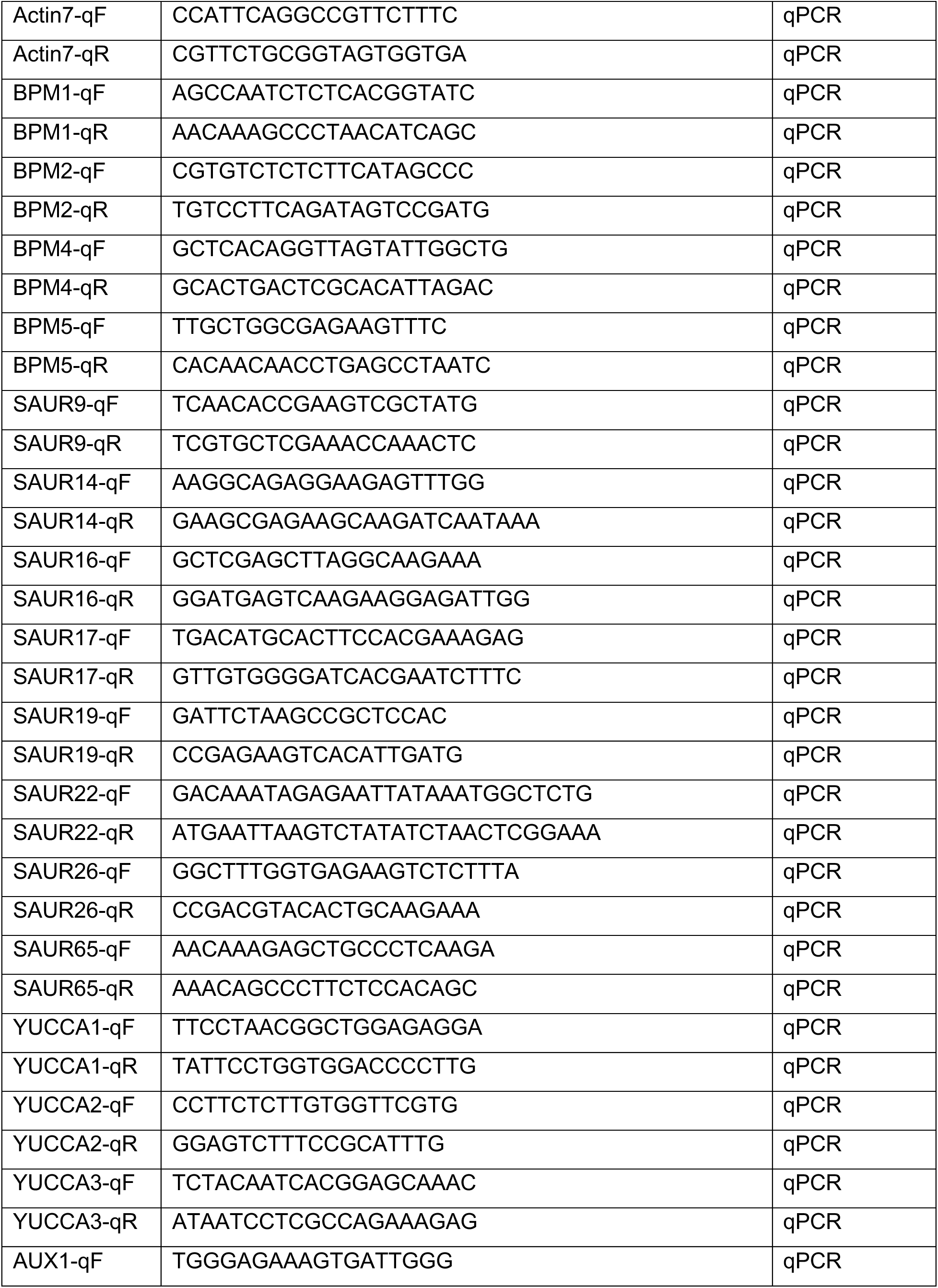
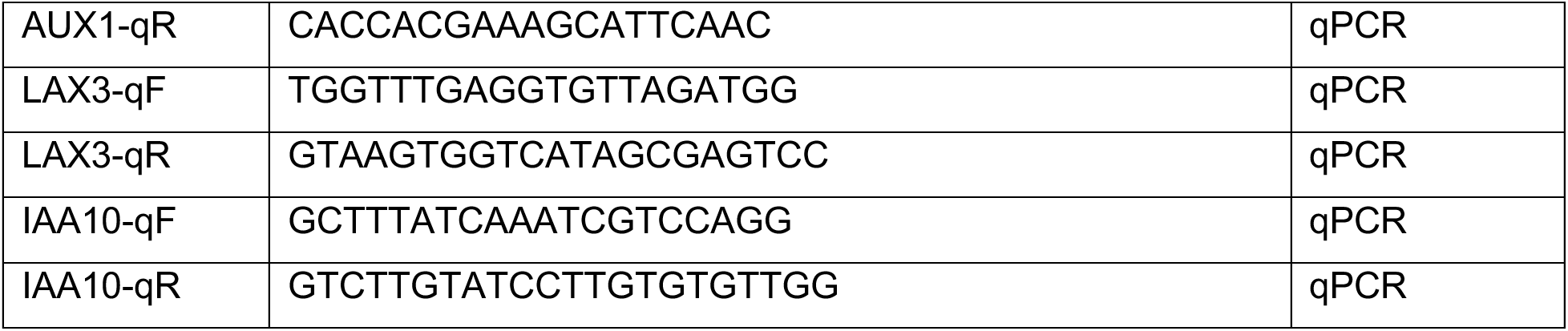
List of primers used in this study.

## References

1. Achard P, Liao L, Jiang C, Desnos T, Bartlett J, Fu X, Harberd NP (2007) DELLAs contribute to plant photomorphogenesis. Plant Physiol 143: 1163–1172

2. Alabadi D, Gil J, Blazquez MA, Garcia-Martinez JL (2004) Gibberellins repress photomorphogenesis in darkness. Plant Physiol 134: 1050–1057

3. Arabidopsis Interactome Mapping C (2011) Evidence for network evolution in an Arabidopsis interactome map. Science 333: 601–607

4. Ban Z, Estelle M (2021) CUL3 E3 ligases in plant development and environmental response. Nat Plants 7: 6–16

5. Calderon Villalobos LI, Lee S, De Oliveira C, Ivetac A, Brandt W, Armitage L, Sheard LB, Tan X, Parry G, Mao H, Zheng N, Napier R, Kepinski S, Estelle M (2012) A combinatorial TIR1/AFB-Aux/IAA co-receptor system for differential sensing of auxin. Nat Chem Biol 8: 477–485

6. Cao M, Chen R, Li P, Yu Y, Zheng R, Ge D, Zheng W, Wang X, Gu Y, Gelova Z, Friml J, Zhang H, Liu R, He J, Xu T (2019) TMK1-mediated auxin signalling regulates differential growth of the apical hook. Nature 568: 240–243

7. Chapman EJ, Greenham K, Castillejo C, Sartor R, Bialy A, Sun TP, Estelle M (2012) Hypocotyl Transcriptome Reveals Auxin Regulation of Growth-Promoting Genes through GA-Dependent and -Independent Pathways. Plos One 7

8. Chen H, Li L, Zou M, Qi L, Friml J (2023) Distinct functions of TIR1 and AFB1 receptors in auxin signaling. Mol Plant 16: 1117–1119

9. Chen L, Bernhardt A, Lee J, Hellmann H (2015) Identification of Arabidopsis MYB56 as a novel substrate for CRL3(BPM) E3 ligases. Mol Plant 8: 242–250

10. Chen L, Lee JH, Weber H, Tohge T, Witt S, Roje S, Fernie AR, Hellmann H (2013) Arabidopsis BPM proteins function as substrate adaptors to a cullin3-based E3 ligase to affect fatty acid metabolism in plants. Plant Cell 25: 2253–2264

11. Chico JM, Lechner E, Fernandez-Barbero G, Canibano E, Garcia-Casado G, Franco-Zorrilla JM, Hammann P, Zamarreno AM, Garcia-Mina JM, Rubio V, Genschik P, Solano R (2020) CUL3(BPM) E3 ubiquitin ligases regulate MYC2, MYC3, and MYC4 stability and JA responses. Proc Natl Acad Sci U S A 117: 6205–6215

12. Clough SJ, Bent AF (1998) Floral dip: a simplified method for Agrobacterium-mediated transformation of Arabidopsis thaliana. Plant J 16: 735–743

13. Dong J, Sun N, Yang J, Deng Z, Lan J, Qin G, He H, Deng XW, Irish VF, Chen H, Wei N (2019) The Transcription Factors TCP4 and PIF3 Antagonistically Regulate Organ-Specific Light Induction of SAUR Genes to Modulate Cotyledon Opening during De-Etiolation in Arabidopsis. Plant Cell 31: 1155–1170

14. Du M, Bou Daher F, Liu Y, Steward A, Tillmann M, Zhang X, Wong JH, Ren H, Cohen JD, Li C, Gray WM (2022) Biphasic control of cell expansion by auxin coordinates etiolated seedling development. Sci Adv 8: eabj1570

15. Du M, Spalding EP, Gray WM (2020) Rapid Auxin-Mediated Cell Expansion. Annu Rev Plant Biol 71: 379–402

16. Dubey SM, Han S, Stutzman N, Prigge MJ, Medvecka E, Platre MP, Busch W, Fendrych M, Estelle M (2023) The AFB1 auxin receptor controls the cytoplasmic auxin response pathway in Arabidopsis thaliana. Mol Plant 16: 1120–1130

17. Fendrych M, Leung J, Friml J (2016) TIR1/AFB-Aux/IAA auxin perception mediates rapid cell wall acidification and growth of Arabidopsis hypocotyls. Elife 5

18. Feng S, Martinez C, Gusmaroli G, Wang Y, Zhou J, Wang F, Chen L, Yu L, Iglesias-Pedraz JM, Kircher S, Schafer E, Fu X, Fan LM, Deng XW (2008) Coordinated regulation of Arabidopsis thaliana development by light and gibberellins. Nature 451: 475–479

19. Gao X, Chen J, Dai X, Zhang D, Zhao Y (2016) An Effective Strategy for Reliably Isolating Heritable and Cas9-Free Arabidopsis Mutants Generated by CRISPR/Cas9-Mediated Genome Editing. Plant Physiol 171: 1794–1800

20. Gendreau E, Traas J, Desnos T, Grandjean O, Caboche M, Hofte H (1997) Cellular basis of hypocotyl growth in Arabidopsis thaliana. Plant Physiol 114: 295–305

21. Genschik P, Sumara I, Lechner E (2013) The emerging family of CULLIN3-RING ubiquitin ligases (CRL3s): cellular functions and disease implications. EMBO J 32: 2307–2320

22. Julian J, Coego A, Lozano-Juste J, Lechner E, Wu Q, Zhang X, Merilo E, Belda-Palazon B, Park SY, Cutler SR, An C, Genschik P, Rodriguez PL (2019) The MATH-BTB BPM3 and BPM5 subunits of Cullin3-RING E3 ubiquitin ligases target PP2CA and other clade A PP2Cs for degradation. Proc Natl Acad Sci U S A 116: 15725–15734

23. Lavy M, Estelle M (2016) Mechanisms of auxin signaling. Development 143: 3226–3229

24. Lechner E, Leonhardt N, Eisler H, Parmentier Y, Alioua M, Jacquet H, Leung J, Genschik P (2011) MATH/BTB CRL3 receptors target the homeodomain-leucine zipper ATHB6 to modulate abscisic acid signaling. Dev Cell 21: 1116–1128

25. Leivar P, Monte E, Oka Y, Liu T, Carle C, Castillon A, Huq E, Quail PH (2008) Multiple phytochrome-interacting bHLH transcription factors repress premature seedling photomorphogenesis in darkness. Curr Biol 18: 1815–1823

26. Li QF, He JX (2016) BZR1 Interacts with HY5 to Mediate Brassinosteroid- and Light-Regulated Cotyledon Opening in Arabidopsis in Darkness. Mol Plant 9: 113–125

27. Li Y, Varala K, Hudson ME (2014) A survey of the small RNA population during far-red light-induced apical hook opening. Front Plant Sci 5: 156

28. Malamy JE, Benfey PN (1997) Organization and cell differentiation in lateral roots of Arabidopsis thaliana. Development 124: 33–44

29. Mooney S, Al-Saharin R, Choi CM, Tucker K, Beathard C, Hellmann HA (2019) Characterization of Brassica rapa RAP2.4-Related Proteins in Stress Response and as CUL3-Dependent E3 Ligase Substrates. Cells 8

30. Morimoto K, Ohama N, Kidokoro S, Mizoi J, Takahashi F, Todaka D, Mogami J, Sato H, Qin F, Kim JS, Fukao Y, Fujiwara M, Shinozaki K, Yamaguchi-Shinozaki K (2017) BPM-CUL3 E3 ligase modulates thermotolerance by facilitating negative regulatory domain-mediated degradation of DREB2A in Arabidopsis. Proc Natl Acad Sci U S A 114: E8528–E8536

31. Ni W, Xu SL, Gonzalez-Grandio E, Chalkley RJ, Huhmer AFR, Burlingame AL, Wang ZY, Quail PH (2017) PPKs mediate direct signal transfer from phytochrome photoreceptors to transcription factor PIF3. Nat Commun 8: 15236

32. Novak O, Henykova E, Sairanen I, Kowalczyk M, Pospisil T, Ljung K (2012) Tissue-specific profiling of the Arabidopsis thaliana auxin metabolome. Plant J 72: 523–536

33. Oh E, Zhu JY, Bai MY, Arenhart RA, Sun Y, Wang ZY (2014) Cell elongation is regulated through a central circuit of interacting transcription factors in the Arabidopsis hypocotyl. Elife 3

34. Prigge MJ, Platre M, Kadakia N, Zhang Y, Greenham K, Szutu W, Pandey BK, Bhosale RA, Bennett MJ, Busch W, Estelle M (2020) Genetic analysis of the Arabidopsis TIR1/AFB auxin receptors reveals both overlapping and specialized functions. Elife 9

35. Ravindran N, Ramachandran H, Job N, Yadav A, Vaishak KP, Datta S (2021) B-box protein BBX32 integrates light and brassinosteroid signals to inhibit cotyledon opening. Plant Physiol 187: 446–461

36. Salehin M, Bagchi R, Estelle M (2015) SCFTIR1/AFB-based auxin perception: mechanism and role in plant growth and development. Plant Cell 27: 9–19

37. Serre NBC, Kralík D, Yun P, Slouka Z, Shabala S, Fendrych M (2021) AFB1 controls rapid auxin signalling through membrane depolarization in root. Nature Plants 7: 1229-+

38. Shin J, Kim K, Kang H, Zulfugarov IS, Bae G, Lee CH, Lee D, Choi G (2009) Phytochromes promote seedling light responses by inhibiting four negatively-acting phytochrome-interacting factors. Proc Natl Acad Sci U S A 106: 7660–7665

39. Skiljaica A, Lechner E, Jagic M, Majsec K, Malenica N, Genschik P, Bauer N (2020) The protein turnover of Arabidopsis BPM1 is involved in regulation of flowering time and abiotic stress response. Plant Mol Biol 102: 359–372

40. Vandenbussche F, Petrasek J, Zadnikova P, Hoyerova K, Pesek B, Raz V, Swarup R, Bennett M, Zazimalova E, Benkova E, Van Der Straeten D (2010) The auxin influx carriers AUX1 and LAX3 are involved in auxin-ethylene interactions during apical hook development in Arabidopsis thaliana seedlings. Development 137: 597–606

41. Vernoux T, Brunoud G, Farcot E, Morin V, Van den Daele H, Legrand J, Oliva M, Das P, Larrieu A, Wells D, Guedon Y, Armitage L, Picard F, Guyomarc’h S, Cellier C, Parry G, Koumproglou R, Doonan JH, Estelle M, Godin C, Kepinski S, Bennett M, De Veylder L, Traas J (2011) The auxin signalling network translates dynamic input into robust patterning at the shoot apex. Mol Syst Biol 7: 508

42. Weber H, Hellmann H (2009) Arabidopsis thaliana BTB/ POZ-MATH proteins interact with members of the ERF/AP2 transcription factor family. FEBS J 276: 6624–6635

43. Xiong Q, Ma B, Lu X, Huang YH, He SJ, Yang C, Yin CC, Zhao H, Zhou Y, Zhang WK, Wang WS, Li ZK, Chen SY, Zhang JS (2017) Ethylene-Inhibited Jasmonic Acid Biosynthesis Promotes Mesocotyl/Coleoptile Elongation of Etiolated Rice Seedlings. Plant Cell 29: 1053–1072

44. Yoo SD, Cho YH, Sheen J (2007) Arabidopsis mesophyll protoplasts: a versatile cell system for transient gene expression analysis. Nat Protoc 2: 1565–1572

45. Zadnikova P, Petrasek J, Marhavy P, Raz V, Vandenbussche F, Ding Z, Schwarzerova K, Morita MT, Tasaka M, Hejatko J, Van Der Straeten D, Friml J, Benkova E (2010) Role of PIN-mediated auxin efflux in apical hook development of Arabidopsis thaliana. Development 137: 607–617

46. Zhao Y, Christensen SK, Fankhauser C, Cashman JR, Cohen JD, Weigel D, Chory J (2001) A role for flavin monooxygenase-like enzymes in auxin biosynthesis. Science 291: 306–309

47. Zheng Y, Cui X, Su L, Fang S, Chu J, Gong Q, Yang J, Zhu Z (2017) Jasmonate inhibits COP1 activity to suppress hypocotyl elongation and promote cotyledon opening in etiolated Arabidopsis seedlings. Plant J 90: 1144–1155

48. Zhong S, Shi H, Xue C, Wei N, Guo H, Deng XW (2014) Ethylene-orchestrated circuitry coordinates a seedling’s response to soil cover and etiolated growth. Proc Natl Acad Sci U S A 111: 3913–3920

49. Zhuang M, Calabrese MF, Liu J, Waddell MB, Nourse A, Hammel M, Miller DJ, Walden H, Duda DM, Seyedin SN, Hoggard T, Harper JW, White KP, Schulman BA (2009) Structures of SPOP-substrate complexes: insights into molecular architectures of BTB-Cul3 ubiquitin ligases. Mol Cell 36: 39–50

